# Covalent Destabilizing Degrader of AR and AR-V7 in Androgen-Independent Prostate Cancer Cells

**DOI:** 10.1101/2025.02.12.637117

**Authors:** Charlotte M. Zammit, Cory M. Nadel, Ying Lin, Sajjan Koirala, Patrick Ryan Potts, Daniel K. Nomura

## Abstract

Androgen-independent prostate cancers, correlated with heightened aggressiveness and poor prognosis, are caused by mutations or deletions in the androgen receptor (AR) or expression of truncated variants of AR that are constitutively activated. Currently, drugs and drug candidates against AR target the steroid-binding domain to antagonize or degrade AR. However, these compounds cannot therapeutically access largely intrinsically disordered truncated splice variants of AR, such as AR-V7, that only possess the DNA binding domain and are missing the ligand binding domain. Targeting intrinsically disordered regions within transcription factors has remained challenging and is considered “undruggable”. Herein, we leveraged a cysteine-reactive covalent ligand library in a cellular screen to identify degraders of AR and AR-V7 in androgen-independent prostate cancer cells. We identified a covalent compound EN1441 that selectively degrades AR and AR-V7 in a proteasome-dependent manner through direct covalent targeting of an intrinsically disordered cysteine C125 in AR and AR-V7. EN1441 causes significant and selective destabilization of AR and AR-V7, leading to aggregation of AR/AR-V7 and subsequent proteasome-mediated degradation. Consistent with targeting both AR and AR-V7, we find that EN1441 completely inhibits total AR transcriptional activity in androgen-independent prostate cancer cells expressing both AR and AR-V7 compared to AR antagonists or degraders that only target the ligand binding domain of full-length AR, such as enzalutamide and ARV-110. Our results put forth a pathfinder molecule EN1441 that targets an intrinsically disordered cysteine within AR to destabilize, degrade, and inhibit both AR and AR-V7 in androgen-independent prostate cancer cells and highlights the utility of covalent ligand discovery approaches in directly targeting, destabilizing, inhibiting, and degrading classically undruggable transcription factor targets.

## Introduction

Prostate cancer remains one of the leading causes of cancer-related deaths in men worldwide, with advanced stages presenting significant therapeutic challenges ^1,2^. Androgen receptor (AR) signaling plays a central role in prostate cancer progression, and therapies targeting the androgen-AR axis, such as androgen deprivation therapy and AR antagonists, are the standard of care for advanced disease ^1,2^. However, many patients eventually develop resistance, leading to androgen-independent prostate cancers. One key driver of resistance is the emergence of truncated AR variants, particularly AR-V7, which lacks the ligand-binding domain, is largely intrinsically disordered and unstructured, and remains constitutively active, thereby bypassing the need for androgen binding ^1–4^. The AR-V7 truncation variants are therapeutically resistant to current clinically approved AR antagonists that target the ligand binding domain such as enzalutamide or AR Proteolysis Targeting Chimeras (PROTACs) like ARV-110 that are in clinical development ^5,6^. Unfortunately, despite the importance of AR-V7 as a drug target for androgen-resistant prostate cancers, the remaining DNA binding domain of AR-V7 is considered “undruggable” by traditional drug discovery approaches because much of AR-V7 is highly intrinsically disordered and structurally poorly defined ^3,4^.

Covalent chemoproteomic platforms have emerged as a powerful approach for targeting previously intractable disease targets that may not possess well-defined binding pockets ^7,8^. Covalent small molecules have been discovered against previously challenging targets, including GTPases, such as KRAS G12C, E3 ubiquitin ligases, splicing factors, RNA binding proteins, and transcription factors ^8–26^. We and others have also successfully targeted intrinsically disordered regions within proteins with covalent ligands to modulate protein function^14,17,18,22,27^.

Targeted protein degradation (TPD) has also arisen as a powerful therapeutic modality for tackling undruggable protein targets by using small molecules to induce the proximity of target proteins with E3 ubiquitin ligases to ubiquitinate and degrade neo-substrate proteins through the proteasome ^28–33^. Another class of degraders has been characterized by their ability to destabilize proteins ^17,18,34–38^. Initial examples of these destabilizers included “hydrophobic tagging” strategies developed by Craig Crews and co-workers that linked greasy adamantane moieties to protein-targeting ligands to distort protein folding and induce degradation through quality control mechanisms ^36–38^. We have also previously identified covalent ligands that target intrinsically disordered cysteines within oncogenic transcription factors such as MYC and CTNNB1 to destabilize and degrade these proteins in a ubiquitin-proteasome-dependent manner ^17,18^. Studies have shown that this type of destabilization-mediated degradation is not just restricted to covalent molecules but also more broadly to reversible inhibitors such as kinase inhibitors ^35^.

In this study, we aimed to identify covalent small molecules that could directly target, destabilize, inhibit, and degrade both AR and AR-V7. Through screening a library of cysteine-reactive electrophiles in a target-based cellular screen, we have identified such a molecule that targets an intrinsically disordered cysteine within the DNA binding domain of both AR and AR-V7 leading to the destabilization and inhibition of AR transcriptional activity followed by subsequent degradation.

## Results

### Cellular Covalent Ligand Screening to Discover AR-V7 Degraders

We sought to identify a cysteine-reactive covalent small-molecule that could degrade AR-V7, the intractable truncation variant of AR in androgen-resistant prostate cancer cells. We screened a library of 1281 cysteine-reactive electrophiles consisting of acrylamides and chloroacetamides to identify molecules that would lower the endogenous levels of AR-V7 specifically tagged with HiBiT in 22Rv1 cells, wherein AR-V7-HiBiT could be detected upon addition of LgBiT to result in a luminescence signal in high-throughput screens **(Figure 1a; Table S1)** ^39^. 22Rv1 cells are androgen-independent prostate cancer cells expressing full-length wild-type AR and the truncation variant AR-V7 ^40,41^. We identified 11 hits that lowered AR-V7-HiBiT by greater than 80%, in which the acrylamide EN1441 was the top hit **(Figure 1a-1b).** EN1441 significantly lowered AR-V7-HiBiT levels, and this reduction was attenuated by pre-treating cells with the proteasome inhibitor bortezomib, demonstrating proteasome-dependent degradation of AR-V7 **(Figure 1c)**. We further validated these results, showing that EN1441 significantly lowered both full-length wild-type AR and untagged AR-V7 in parental 22Rv1 cells in a proteasome-dependent manner **(Figure 1d-1e).** Interestingly, EN1441-mediated loss of AR and AR-V7 was not attenuated by the neddylation inhibitor MLN4924 that blocks the activity of the Cullin family of E3 ubiquitin ligases, indicating that this degradation was not occurring through a Cullin E3 ligase **(Figure S1a-S1b)**^42^. EN1441 showed dose-responsive degradation of AR and AR-V7 with rapid loss occurring at one hour of treatment **(Figure 1f-1g)**. This loss of AR was not only observed in 22Rv1 cells but also androgen-dependent LNCaP prostate cancer cells **(Figure S1c).** We also showed that the covalent acrylamide warhead was necessary since the unreactive propenamide analog CMZ139 did not show a loss of AR and AR-V7 when compared to EN1441 **(Figure 1h-1i).** Despite the modest potency of EN1441, quantitative proteomic profiling of EN1441 showed selective degradation of AR and AR-V7 in 22Rv1 cells **(Figure 1j, Table S2).** Note that by proteomics, given the sequence identity in the DNA binding region between AR and AR-V7, we cannot distinguish between them.

**Figure 1.**
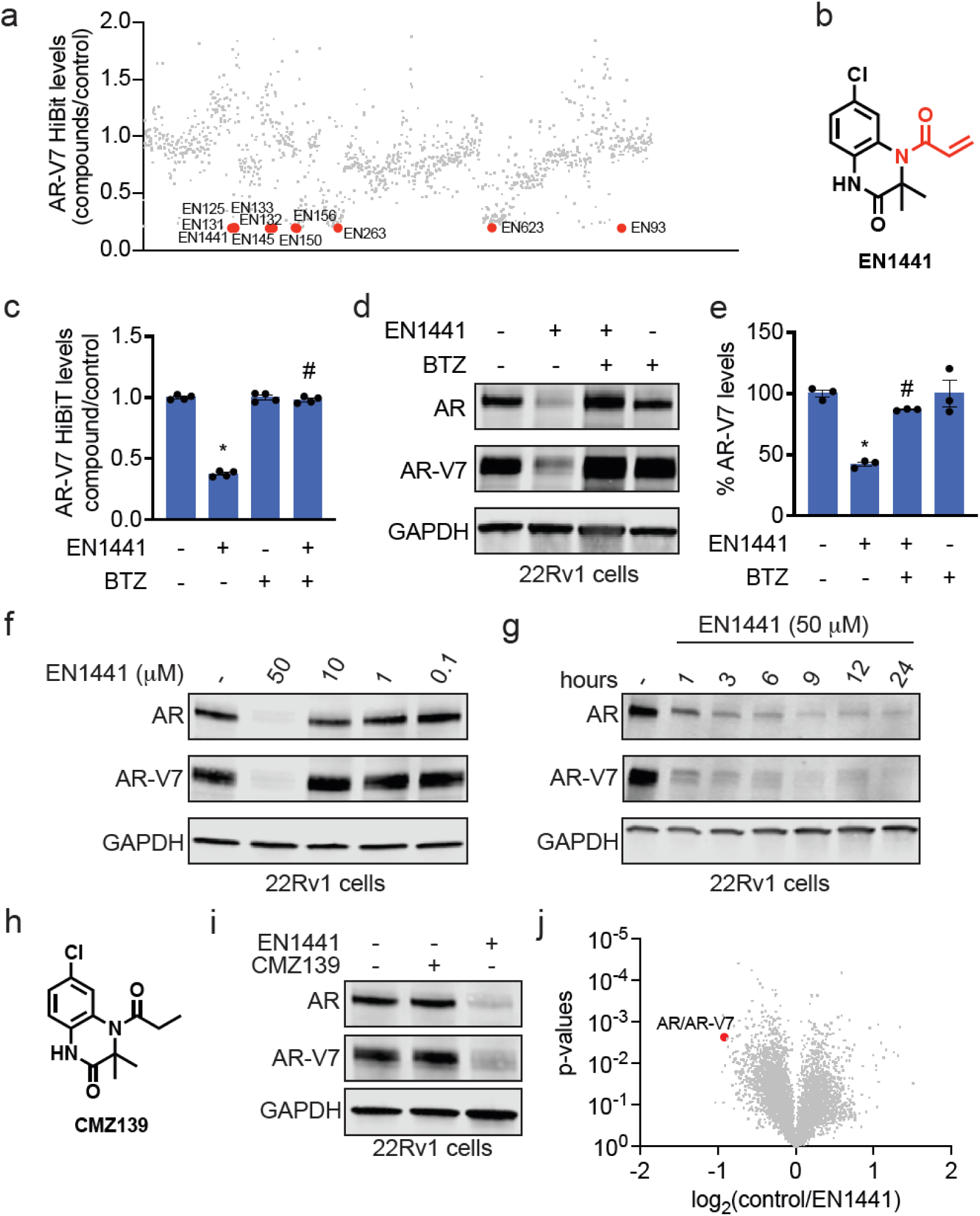
Cellular covalent ligand screening to discover AR-V7 degraders. **(a)** Covalent ligand screen in 22Rv1 cells expressing an endogenously C-terminally tagged HiBiT peptide on AR-V7. These cells were treated with DMSO vehicle or cysteine-reactive covalent ligand (50 μM) for 24 h, after which HiBiT-AR-V7 was detected by luminescence through detection with LgBiT. Individual compounds, structures, and data points are shown in **Table S1.** Data is shown as the ratio of luminescence between compound versus vehicle treatment. Highlighted in red are compounds that showed >80 % reduction in AR-V7-HiBiT. Data are from n=1 biological replicate/group. **(b)** The structure of the top hit EN1441 with cysteine-reactive acrylamide warhead highlighted in red. **(c)** Proteasome-dependence of EN1441-mediated AR-V7-HiBiT loss. 22Rv1 cells expressing AR-V7- HiBiT were pre-treated with DMSO vehicle or bortezomib (BTZ) for 1 h prior to treatment of cells with DMSO vehicle or EN1441 (50 μM) for 24 h, after which AR-V7-HiBiT levels were detected by LgBiT and resulting luminescence. Data were normalized to respective cell viability for each group. **(d)** AR and AR-V7 levels in 22Rv1 cells treated with EN1441. 22Rv1 parental cells were pre-treated with DMSO vehicle or BTZ (1 μM) for 1 h prior to treatment of cells with DMSO vehicle or EN1441 (50 μM) for 24 h, and AR and AR-V7 levels and loading control GAPDH were detected by Western blotting. **(e)** Quantification of experiment in **(d)**. **(f)** Dose-response of AR and AR-V7 degradation with EN1441 treatment. 22Rv1 cells were treated with DMSO or EN1441 for 24 h, and AR, AR-V7, and GAPDH levels were detected by Western blotting. **(g)** Time-course of AR and AR-V7 degradation with EN1441 treatment. 22Rv1 cells were treated with DMSO or EN1441 (50 μM), and AR, AR-V7, and GAPDH levels were detected by Western blotting. **(h)** Structure of unreactive analog of EN1441, CMZ139. **(i)** AR and AR-V7 levels with CMZ139 treatment. 22Rv1 cells were treated with EN1441 or CMZ139 (50 μM) for 16 h, and AR, AR-V7, and GAPDH levels were detected by Western blotting. **(j)** Quantitative tandem mass tagging (TMT)-based proteomic profiling of EN1441. 22Rv1 cells were treated with DMSO vehicle or EN1441 (50 μM) for 16 h and resulting cell lysates were subjected to quantitative proteomic profiling. Shown in red is AR/AR-V7. Full data can be found in **Table S2.** Data in **(c)** are from n=4, and data in **(c-g, i, j)** are from n=3 biologically independent replicates per group. Blots in **(d,f,g,i)** are representative. Bar graphs in **(c,e)** show individual replicate values and average ± sem. Significance in **(c,d)** expressed as *p<0.05 compared to vehicle-treated controls and #p<0.05 compared to EN1441-treated groups.

### EN1441 Directly Engages AR and AR-V7

To determine whether EN1441 may be acting through direct interactions with AR-V7, we performed a gel-based activity-based protein profiling (ABPP) experiment competing EN1441 against labeling of pure human AR-V7 protein with a rhodamine-conjugated cysteine-reactive iodoacetamide probe ^23,43,44^. EN1441 showed dose-responsive binding to AR-V7 *in vitro* **(Figure 2a).** Analysis of the EN1441 site of modification on AR-V7 by mass spectrometry-based detection of the EN1441 adduct on pure AR-V7 protein revealed cysteine 125 (C125) as the primary site of modification **(Figure 2b)**. Interestingly, C125 is within a predicted intrinsically disordered region of AR-V7 based on PONDR^®^ prediction **(Figure 2b; Figure S1d)** ^45^. This C125 is conserved in both full-length AR and AR-V7, explaining why we observe degradation of both AR and AR-V7.

**Figure 2.**
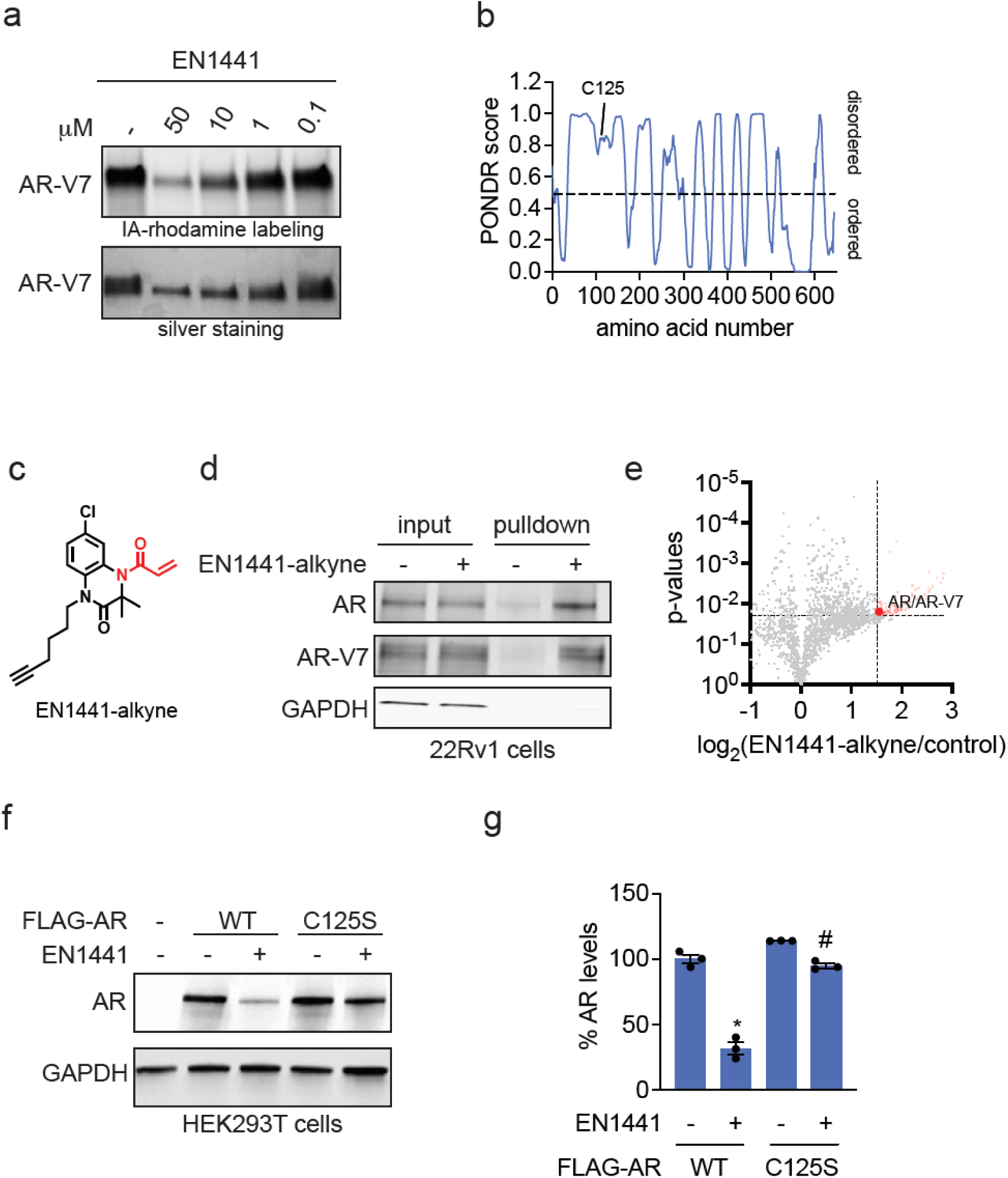
EN1441 directly engages AR and AR-V7. **(a)** Gel-based ABPP analysis of EN1441 with pure AR- V7 protein. AR-V7 pure protein was pre-incubated with DMSO or EN1441 for 30 min prior to labeling with rhodamine-conjugated iodoacetamide (IA-rhodamine) for 1 h after which proteins were separated by SDS/PAGE and visualized by in-gel fluorescence, and loading was assessed by silver staining. **(b)** EN1441 modifies intrinsically disordered C125 on AR-V7. PONDR^®^ prediction of AR-V7 sequence predicting disordered (above 0.5) or ordered region (below 0.5), and highlighting C125 site of modification based on labeling of EN1441 (50 μM, 1 h) with pure AR-V7 protein and analyzing resulting tryptic digests by LC-MS/MS. Data from this experiment is in **Figure S1d. (c)** Structure of EN1441-alkyne probe. **(d)** AR and AR-V7 enrichment with EN1441-alkyne probe. 22Rv1 cells were pre-treated with BTZ (1 μM) for 1 h prior to treatment of cells with DMSO vehicle or EN1441-alkyne (5 μM, 4.5 h). Resulting lysates were subjected to copper-catalyzed azide-alkyne cycloaddition (CuAAC) with an azide-functionalized biotin handle, and probe-modified proteins were avidin-enriched, eluted, separated by SDS/PAGE and AR, AR-V7, and unrelated control GAPDH were detected by Western blotting. Shown are protein levels from input and pulldown proteome. **(e)** EN1441-alkyne pulldown chemoproteomic profiling. 22Rv1 cells were pre-treated with BTZ (1 μM) for 1 h prior to treatment of cells with DMSO vehicle or EN1441-alkyne (5 μM, 4.5 h). Resulting lysates were subjected to CuAAC with an azide-functionalized biotin handle, and probe-modified proteins were avidin-enriched, eluted, and proteins were analyzed by TMT-based quantitative proteomic profiling. Shown are proteins detected and enrichment ratio between EN1441-alkyne versus vehicle-treated controls. Shown in red are proteins enriched by >log_2_ 1.5 with p<0.02. Highlighted in bold red is AR/AR-V7. Data from this experiment is in **Table S3. (f)** AR wild-type (WT) versus C125S mutant degradation by EN1441. HEK293T cells were transfected with mock or FLAG-tagged WT or C125S mutant full-length AR for 24 h, after which cells were treated with DMSO vehicle or EN1441 (50 μM, 6h), and FLAG (FLAG-AR) and loading control GAPDH levels were assessed by Western blotting. **(g)** Quantification experiment in **(f)**. Data in **(a,d,e,f,g)** are from n=3 biologically independent replicates per group. Gels and blots in **(a,d,f)** are representative. Bar graph in **(g)** shows individual replicate values and average ± sem. Significance in **(g)** expressed as *p<0.05 compared to vehicle-treated FLAG-AR WT expressing cells and #p<0.05 compared to EN1441-treated FLAG-AR WT expressing cells.

To determine AR and AR-V7 engagement and proteome-wide selectivity of EN1441 in cells, we synthesized an alkyne-functionalized probe of EN1441, EN1441-alkyne **(Figure 2c)**. AR and AR-V7, but not unrelated targets such as GAPDH, were enriched by EN1441-alkyne treatment in 22Rv1 cells compared to vehicle-treated control cells **(Figure 2d)**. Chemoproteomic profiling of EN1441-alkyne also showed 2.9-fold significant (p<0.05) enrichment and engagement of AR and AR-V7 compared to vehicle-treated controls **(Figure 2e, Table S3)**. Out of 1830 proteins detected and quantified, we identified another 110 targets that were significantly enriched by EN1441-alkyne **(Figure 2e, Table S3)**. Thus, EN1441 shows modestly selective and direct engagement of AR and AR-V7 in 22Rv1 cells.

Given the various off-targets of EN1441 identified, we sought to confirm on-target activity. EN1441- mediated degradation of wild-type AR was completely attenuated upon mutation of C125 to serine, demonstrating that the observed AR degradation was through direct engagement of C125 **(Figure 2f-2g)**.

### Functional Inhibition of AR Transcriptional Activity

Given the pronounced loss of both AR and AR-V7 in androgen-independent 22Rv1 prostate cancer cells, we next wanted to determine whether EN1441 inhibited AR transcriptional activity in these cells. We also wanted to examine whether EN1441 could more robustly inhibit total AR transcriptional activity in these cells compared to the standard of care for androgen-resistant prostate cancers, such as enzalutamide or AR PROTACs in clinical development, such as ARV110 ^5,6^. EN1441 showed nearly complete inhibition of AR transcriptional luciferase reporter activity in 22Rv1 cells with an EC_50_ of 4.2 μM, which is well below the concentration needed for AR or AR-V7 degradation **(Figure 3a)**. We observed little to no inhibition of AR transcriptional activity with the non-reactive version of EN1441, CMZ139 **(Figure 3b)**. The AR C125S mutant showed resistance to EN1441-mediated inhibition of AR transcriptional reporter activity observed in HEK293T cells expressing wild-type AR **(Figure 3c).** Expectedly, the non-degradative competitive AR antagonist enzalutamide did not degrade AR or AR-V7 and only partially inhibited AR transcriptional activity in 22Rv1 cells with only 44 % inhibition at the highest concentration tested **(Figure 3d-3e).** Also consistent with previous reports, the AR PROTAC ARV110 that also targets the steroid binding domain, selectively degraded only the full-length AR, but not AR-V7, and showed more, 70%, but not complete inhibition of AR transcriptional activity in 22Rv1 cells **(Figure 3d-3e).** EN1441, which degrades both AR and AR-V7, showed a more robust inhibition of total AR transcriptional activity than enzalutamide and ARV110, with 90 % inhibition observed at the highest concentration tested **(Figure 3d-3e)**.

**Figure 3.**
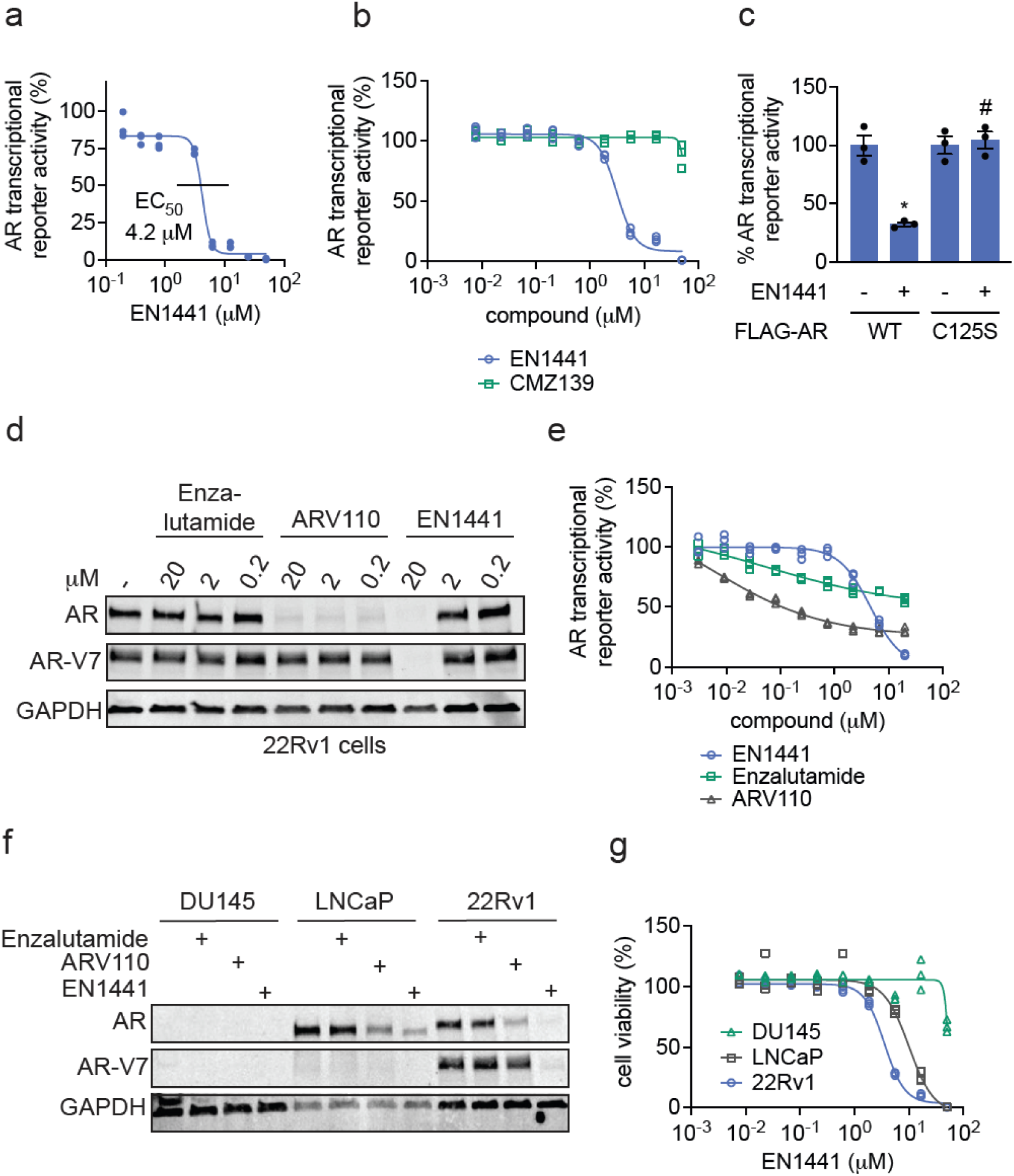
Functional inhibition of AR transcriptional activity. **(a, b)** EN1441 and CMZ139 effects on AR luciferase reporter transcriptional activity. 22Rv1 cells stably expressing an androgen response element (ARE) upstream of a luciferase reporter were treated with DMSO vehicle, EN1441 **(a,b)**, or CMZ139 **(b)** for 24 h, and AR luciferase reporter activity was assessed. The 50 % effective concentration of EN1441 **(a)** was calculated to be 4.2 μM. **(c)** AR luciferase reporter assay in HEK293T cells transiently expressing FLAG-AR WT or FLAG- AR C125S mutant protein and the ARE luciferase reporter co-treated with dihydroxytestosterone (10 nM) and DMSO vehicle or EN1441 (25 μM) for 24 h. AR transcriptional reporter activity was normalized to cell viability as assessed by CellTiter-Glo. **(d)** AR and AR-V7 levels with enzalutamide, ARV110, and EN1441 treatment. 22Rv1 cells were treated with enzalutamide, ARV110, or EN1441 for 24 h, and AR, AR-V7, and loading control GAPDH levels were assessed by Western blotting. **(e)** AR luciferase reporter activity. 22Rv1 cells expressing an ARE luciferase reporter were treated with DMSO vehicle, EN1441, enzalutamide, or ARV110 for 24 h, and AR luciferase reporter activity was assessed. **(f)** AR and AR-V7 levels with enzalutamide, ARV110, and EN1441 treatment. DU145, LNCaP, and 22Rv1 prostate cancer cells were treated with 20 μM enzalutamide, ARV110, or EN1441 for 24 h, and AR, AR-V7, and GAPDH levels were assessed by Western blotting. **(g)** Cell viability with EN1441 treatment in DU145, LNCaP, and 22Rv1 cells. DU145, LNCaP, or 22Rv1 cells were treated with EN1441 for 72 h, after which cell viability was assessed by Cell Titer Glo. Data in **(a-g)** are from n=3 biologically independent replicates per group. Bar graph in **(e)** shows individual replicate values and average ± sem. Blots in **(d,f)** are representative. Plots in **(a,b,e,g)** show individual replicate values.

We next tested whether EN1441 compromises prostate cancer cell viability in a manner correlated with AR expression. EN1441 showed dose-responsive impairments in cell viability from 72-hour treatment in both 22Rv1 and LNCaP cells that express AR or AR-V7, but much less so in androgen-independent DU145 cells that do not express AR **(Figure 3f-3g)**. While these data do not prove that the cell viability impairments observed with EN1441 are only through AR-targeting, we observe differential sensitivity to EN1441 based on AR expression.

### Transcriptomic Profiling of EN1441

Given the near complete inhibition of AR transcriptional reporter activity in cells compared to the AR antagonist enzalutamide or our negative control compound CMZ139, we next sought to determine whether this functional inhibition by EN1441 leads to downregulation of AR-responsive genes in 22Rv1 cells. We performed RNA sequencing comparing transcriptional changes conferred by enzalutamide, EN1441, and CMZ139 **(Figure 4a-4c; Table S4).** Consistent with only partial or no inhibition of AR transcriptional activity with enzalutamide and CMZ139, respectively, we observed only a few significant transcriptional changes with the treatment of 22Rv1 cells with either compound with only four genes significantly down-regulated by >2-fold with enzalutamide treatment and two genes significantly up-regulated by >2-fold with CMZ139 treatment **(Figure 4a, 4c; Table S4)**. In contrast, EN1441 treatment led to significant down-regulation of 921 transcripts and up-regulation of 471 transcripts by >2-fold **(Figure 4b; Table S4)**. Delving into this transcriptomic signature, EN1441 showed significant changes across hallmark androgen-response genes, compared to enzalutamide and CMZ139 **(Figure 4d; Table S4)** ^46–48^. Known target genes such as *NUP210*, *UBE2C*, *NPR3*, *SLC3A*, and *EDN2* were significantly more up-regulated or down-regulated with EN1441 treatment compared to enzalutamide or CMZ139 treatment **(Figure 4e-4i)**. We also quantified the expression of prostate-specific antigen or *KLK3* and showed that EN1441, but not enzalutamide or CMZ139, significantly down-regulated *KLK3* mRNA levels in 22Rv1 cells **(Figure 4j).** These results further indicated that, through complete inhibition of AR transcriptional activity by EN1441, we observed a more extensive effect on downstream transcriptional targets **(Figure 4a-4j)**.

**Figure 4.**
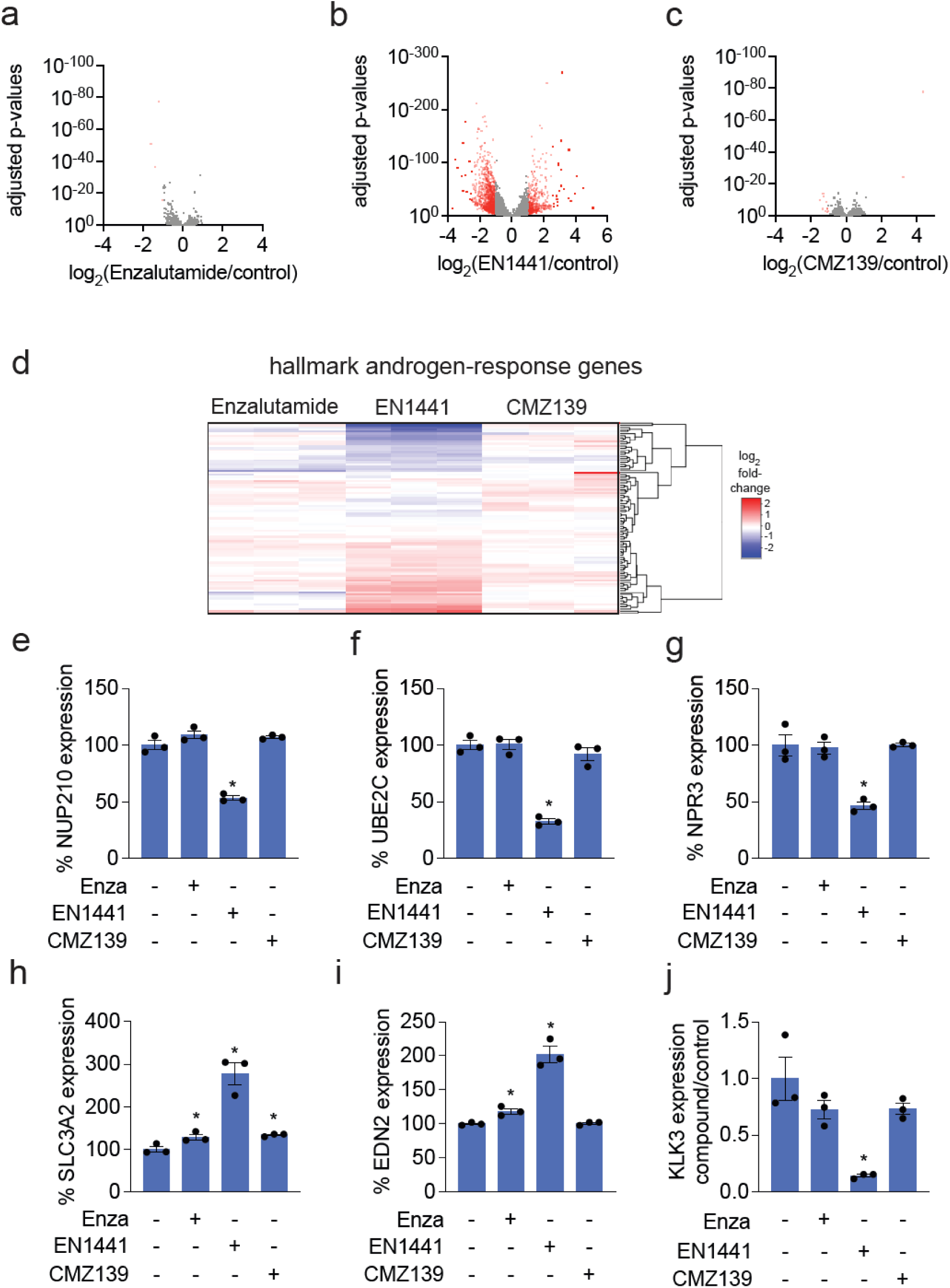
Transcriptomic profiling of EN1441. **(a-c)** Transcriptomic profiling of enzalutamide **(a)**, EN1441 **(b)**, and CMZ139 **(c)** in 22Rv1 cells. 22Rv1 cells were treated with 5 μM enzalutamide, ARV110, or EN1441 for 20 h, after which extracted RNA was subjected to RNA sequencing and transcripts were quantified. Shown in red are significantly changed genes (p<0.05) by >2-fold. **(d)** Hallmark androgen-response genes from the experiment described in **(a-c)** are shown. **(e-i)** Representative androgen response genes from experiment described in **(a-d)** are plotted. **(j)** RT-qPCR analysis of prostate-specific antigen or KLK3 in 22Rv1 cells. 22Rv1 cells were treated with with 5 μM enzalutamide, ARV110, or EN1441 for 20 h, after which extracted RNA was subjected to qPCR analysis. Data shown in **(a-j)** are from n=3 biologically independent replicates per group. Data in **(a-d)** can be found in **Table S4.** Bar graphs in **(e-j)** show individual replicate values and average ± sem. Significance in **(e-j)** expressed as *p<0.05 compared to vehicle-treated controls.

### Understanding Mechanism of AR and AR-V7 Degradation

We next sought further to understand the EN1441-mediated AR and AR-V7 degradation mechanism. Given our previous discoveries with MYC and CTNNB1, where we discovered direct covalently acting small molecules that led to the destabilization and degradation of these proteins, we investigated whether EN1441 acted through similar mechanisms. We performed a thermal denaturation sensitivity assay of EN1441 on both pure AR-V7 protein as well as in 22Rv1 cells **(Figure 5a-5d)** ^49^. EN1441 caused a pronounced and significant destabilization of pure AR-V7 protein as well as a substantial destabilization of both AR and AR-V7 in 22Rv1 cells with over a seven-degree shift in 50 % melting temperature (T_m_) shift for AR-V7 **(Figure 5a-5d)**.

**Figure 5.**
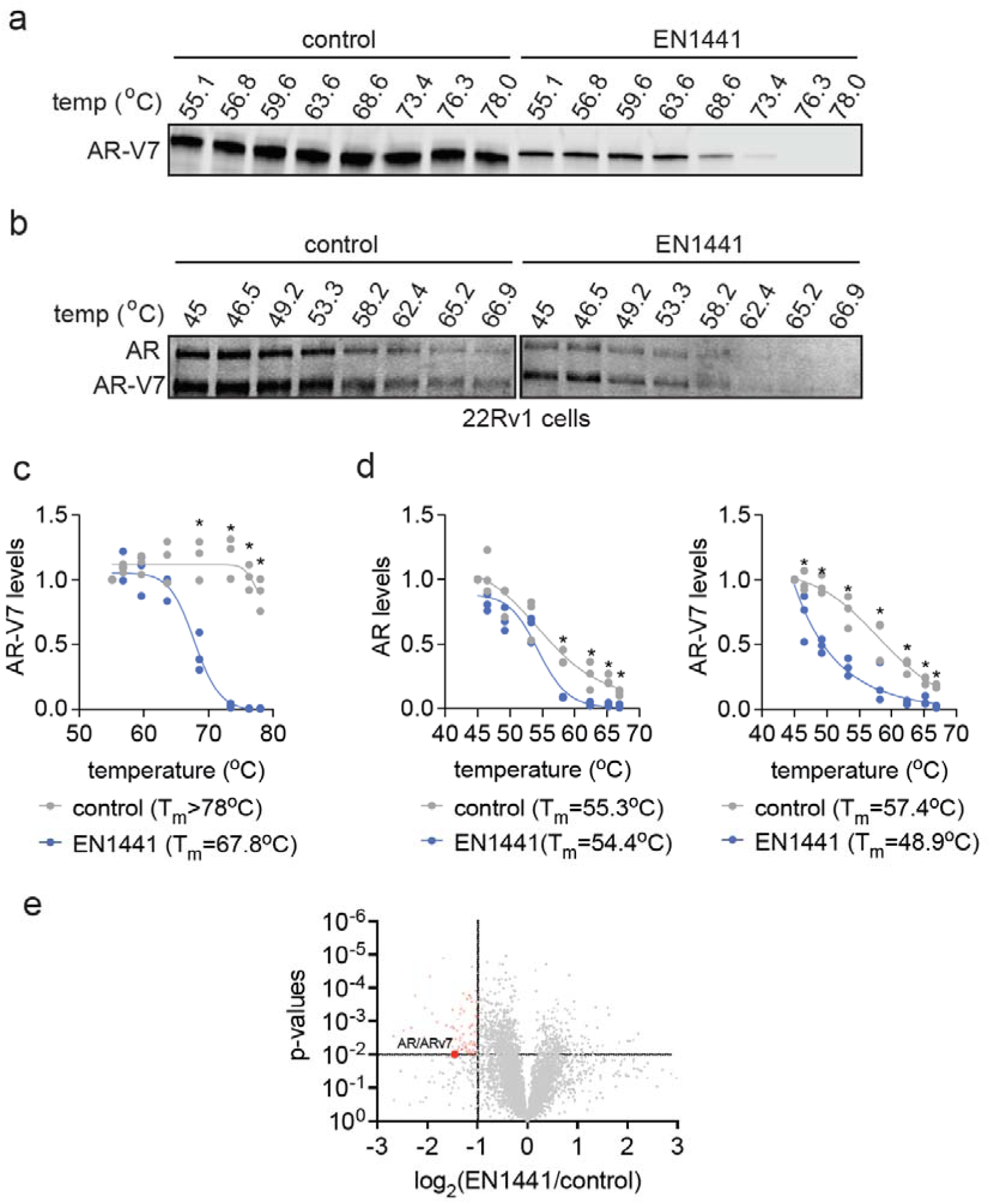
Understanding the mechanism of AR and AR-V7 degradation. **(a)** Thermal denaturation sensitivity assay analysis of EN1441 on pure AR-V7 protein. Pure AR-V7 protein was treated with DMSO vehicle or EN1441 (50 μM) for 1 h, after which the protein was heated to the designated temperatures, insoluble protein was removed, and remaining protein was separated by SDS/PAGE and AR-V7 was detected by Western blotting. **(b)** Thermal denaturation sensitivity assay analysis of EN1441 treatment in 22Rv1 cells. 22Rv1 cells were treated with DMSO or EN1441 (50 μM) for 1 h, after which cells were heated to the designated temperatures, insoluble proteins were removed, and the remaining protein was separated by SDS/PAGE and AR and AR-V7 were detected by Western blotting. **(c)** Quantification of data from **(a)**. **(d)** Quantification of data from **(b)**. **(e)** Thermal denaturation sensitivity assay based-TMT proteomic profiling on EN1441. 22Rv1 cells were treated with DMSO vehicle or EN1441 (50 μM) for 1 h, after which cells were heated over a range of temperatures, insoluble proteins were removed, and remaining proteomes from control temperature groups were combined, treated temperature groups were combined, and subjected to TMT-based quantitative proteomics to identify proteins that were destabilized by EN1441 treatment. Proteins in red were significantly (p<0.01) destabilized by EN1441 by >2-fold. Data from **(a-e)** are from n=3 biologically independent replicates per group. Blots in **(a,b)** are representative. Plots in **(c,d)** show individual replicate values. Data in **(e)** can be found in **Table S5**. Significance is shown as *p<0.05 compared to EN1441-treated groups.

Given the significant change in T_m_, we wanted to confirm that EN1441 was not causing destabilization and denaturation of the whole proteome in a non-specific manner, as that would lead to considerable toxicity. We performed thermal denaturation sensitivity assay-based quantitative proteomic profiling on EN1441 in 22Rv1 cells. We showed relatively selective AR and AR-V7 destabilization with only 85 other proteins that showed significant (p<0.01) destabilization by >2-fold out of 6009 proteins quantified **(Figure 5e; Table S5).**

### Destabilization Versus Degradation of AR and AR-V7 by EN1441

Having shown that EN1441 pronouncedly destabilized AR and AR-V7 in cells, we next sought to determine whether this destabilization led to precipitation or aggregation of AR and AR-V7 in cells and whether this destabilization or aggregation versus degradation was responsible for the inhibition of AR transcriptional activity in cells. To determine whether EN1441 treatment impacts AR and AR-V7 solubility in 22Rv1 cells, we separated cellular extracts from control or EN1441 treated cells into soluble and insoluble fractions and examined AR and AR-V7 abundance. As expected, treatment with EN1441 led to the loss of both soluble and insoluble fractions of AR and AR-V7 in 22Rv1 cell lysate **(Figure 6a)**. However, upon blocking degradation with bortezomib, EN1441-induced a clear repartitioning of AR and AR-V7 to the insoluble fraction compared to vehicle-treated cells **(Figure 6a).** Similar results were obtained if AR and AR-V7 degradation was blocked by the E1 ubiquitin-conjugating enzyme inhibitor TAK243 **(Figure S2a)**^50^. These results indicated that EN1441 engagement of AR and AR-V7 led to the destabilization and aggregation of AR and AR-V7 in a ubiquitin and proteasome-independent manner and that the insoluble versions of these proteins were subsequently eliminated through ubiquitination and subsequent proteasomal degradation. Previous studies have shown that electrophiles can produce general cell stress and non-specific proteotoxic stress, leading to stress granules ^51^. However, we showed that, in contrast to the treatment of cells with a known cellular stressor and stress-granule-inducing sodium arsenite, EN1441 did not show any stress granule formation in cells **(Figure S2b)**. This lack of stress granule formation is consistent with our quantitative proteomic profiling, chemoproteomic profiling, and thermal denaturation sensitivity assay-based proteomic profiling, showing an overall relative selectivity of EN1441 **(Figures 1j, 2e, and 5e)**.

**Figure 6.**
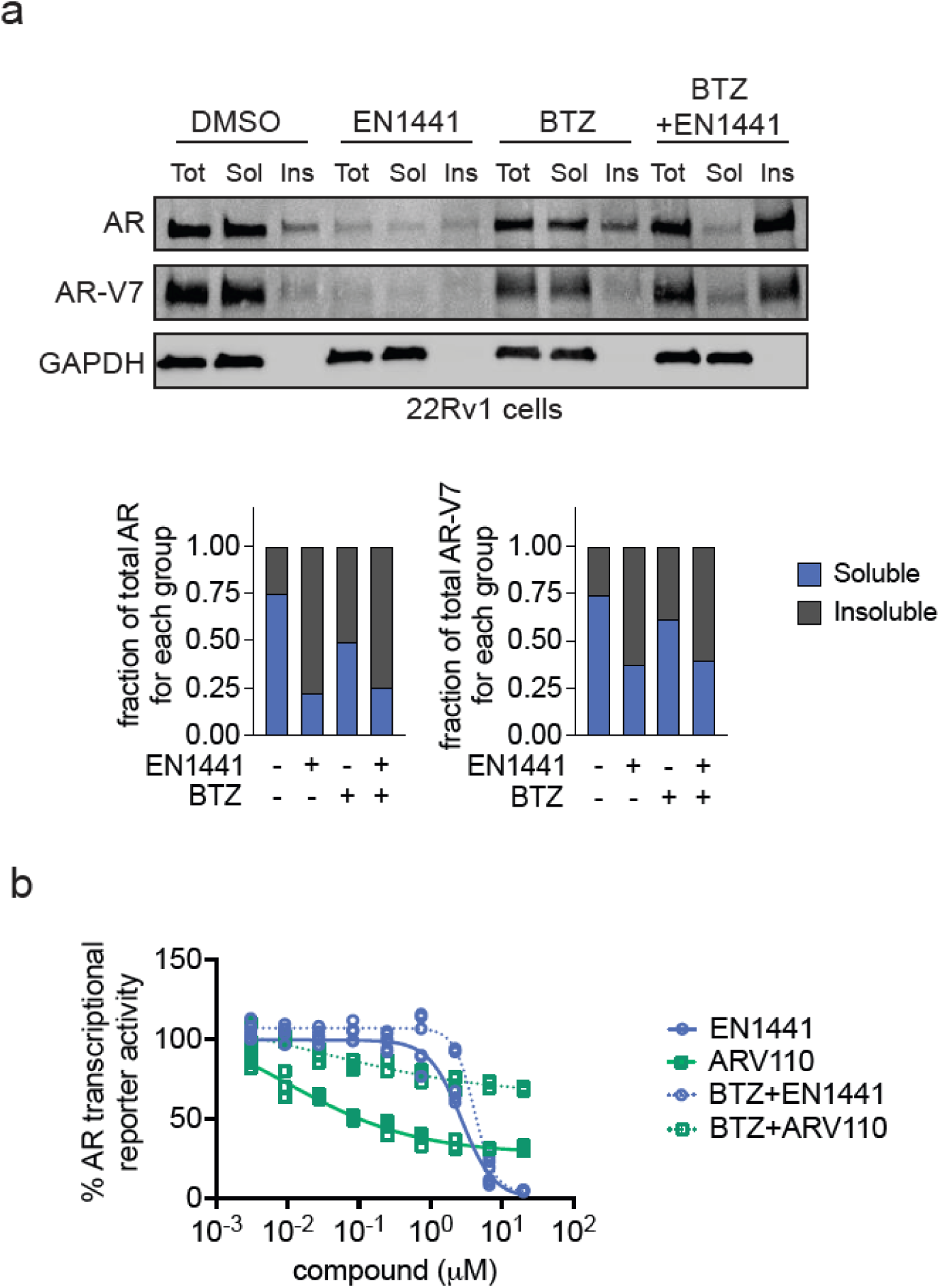
Destabilization versus degradation of AR and AR-V7 by EN1441. **(a)** Total, soluble, and insoluble AR, AR-V7, and loading control GAPDH levels in 22Rv1 cells. 22Rv1 cells were pre-treated with DMSO or bortezomib (1 μM) for 1 h prior to treatment of cells with DMSO vehicle or EN1441 (20 μM) for 24 h. Soluble (sol) and insoluble (ins) fractions were isolated from total cell lysate (tot), separated on SDS/PAGE, and AR, AR-V7, and GAPDH levels were detected by Western blotting. Bar graph below shows quantification of data showing AR and AR-V7 soluble and insoluble levels in relation to total AR in each treatment group. **(b)** AR transcriptional reporter activity in 22Rv1 cells. 22Rv1 cells were pre-treated with DMSO or bortezomib (1 μM) for 1 h prior to treatment of cells with DMSO vehicle or EN1441 for 24 h. AR luciferase reporter activity was subsequently assessed. Data in **(a-b)** are from n=3 biologically independent replicates per group. Blot in **(a)** is representative. Plot in **(b)** shows individual biological replicates.

We next assessed whether the AR transcriptional inhibition observed in 22Rv1 cells was through AR and AR-V7 destabilization or degradation. We convincingly showed that while inhibition of AR transcriptional activity by the AR PROTAC ARV110 showed significant attenuation upon treatment with proteasome inhibitor, EN1441 inhibition of AR activity remained comparable with proteasome inhibitor treatment **(Figure 6b)**. Overall, these results indicated that EN1441 directly engages C125 of AR and AR-V7, leading to destabilization and aggregation of AR and AR-V7 and the subsequent inhibition of AR transcriptional activity and that ubiquitin-mediated proteasomal degradation is a later consequence of destabilization and aggregation.

## Discussion

In this study, we identified a covalent small molecule, EN1441, that selectively targets and degrades both full-length AR and its constitutively active splice variant AR-V7 in androgen-independent prostate cancer cells. Unlike current therapies that target the ligand-binding domain (LBD) of AR, EN1441 covalently modifies a conserved cysteine residue (C125) within the intrinsically disordered DNA-binding domain. This modification leads to the destabilization, aggregation, and subsequent proteasome-dependent degradation of AR and AR- V7. Importantly, EN1441 achieved near-complete inhibition of AR transcriptional activity, surpassing the efficacy of existing AR antagonists like enzalutamide and PROTAC degraders such as ARV-110, which are ineffective against AR variants lacking the ligand binding domain ^3,4^.

The ability of EN1441 to directly engage an intrinsically disordered region challenges the long-held view that such domains within transcription factors are “undruggable”. By covalently modifying C125, EN1441 induces conformational destabilization that not only disrupts AR function, but also targets it for degradation. This approach demonstrates that covalent ligand discovery can effectively expand the druggable proteome to include proteins and domains previously considered intractable due to their lack of well-defined structures. Our findings suggest that targeting intrinsically disordered regions with covalent small molecules may be a viable strategy for modulating the activity of other challenging proteins involved in cancer and other diseases.

Mechanistically, EN1441 mode of action appears to be two-fold: initial destabilization and aggregation of AR and AR-V7, followed by proteasomal degradation. The inhibition of AR transcriptional activity occurs early, concomitant with protein destabilization and independent of degradation, as evidenced by the sustained transcriptional inhibition even when proteasome activity is blocked. This contrasts with the mechanism of PROTACs like ARV-110, which rely on ubiquitination and subsequent proteasomal degradation to inhibit protein function. The aggregation induced by EN1441 may sequester AR and AR-V7 in insoluble complexes, preventing them from engaging in transcriptional regulation, thereby providing a distinct therapeutic mechanism.

While EN1441 shows promise as a lead compound, several considerations warrant further investigation. Although proteomic analyses indicate a degree of selectivity, EN1441 does engage other cellular proteins, which could contribute to off-target effects. EN1441 is also modestly potent, showing mid-micromolar potency. Future work should focus on medicinal chemistry efforts to optimize the specificity and potency of EN1441 to minimize potential side effects. Additionally, further medicinal chemistry efforts are required to optimize metabolic stability and oral bioavailability to enable *in vivo* studies to assess the therapeutic efficacy of these classes of compounds in animal models of androgen-independent prostate cancer. Understanding the long-term effects of AR and AR-V7 destabilization and aggregation on cellular homeostasis will also be crucial for developing safe and effective therapies. As such, EN1441 should be considered a proof-of-concept “pathfinder molecule” that provides a potential path for directly targeting an intractable therapeutic target such as AR-V7 ^52^.

In summary, our study introduces a provocative strategy for targeting intrinsically disordered regions within transcription factors using covalent ligands. EN1441 is a proof-of-concept molecule that effectively destabilizes, inhibits, and degrades both AR and AR-V7. It offers a potential therapeutic avenue for combating androgen-independent prostate cancers resistant to current AR-targeted therapies.

## Acknowledgment

We thank the members of the Nomura Research Group and Amgen for critically reading the manuscript. This work was supported by Amgen Inc., the National Science Foundation Molecular Foundations for Biotechnology (MFB) grant, the UC Berkeley Molecular Therapeutics Initiative (MTI), the Mark Foundation for Cancer Research ASPIRE Award, Bakar Fellows Award, and the National Institutes of Health (R35CA263814, R01CA240981). We also thank Hasan, Lund, and the UC Berkeley NMR facility in the College of Chemistry (CoC-NMR) for spectroscopic assistance. Instruments in the College of Chemistry NMR facility are partly supported by NIH S10OD024998.

## Author Contributions

DKN, PRP, CMZ, CMN co-conceived the project. CMZ, CMN, YL, SK, DKN, and PRP designed and interpreted experiments. CMZ, CMN, YL, and DKN performed experiments. CMZ, CMN, DKN, and PRP wrote the manuscript.

## Methods

### Materials

The cysteine-reactive covalent ligand library was purchased from Enamine LLC. The synthesis and characterization of the non-reactive analog CMZ139 and the alkyne probe EN1441-alkyne are described in the Supporting Information. EN1441 was purchased from Enamine (cat. #Z2738287108). The compound structures are all listed in **Table S1**.

### Equation for Determining EC_50_ Values

The following equation was used for curve-fitting based on the experimental data and calculating EC_50_ values *Y* = bottom + (top − bottom)/(1 + EC_50_/*X*) ^hill slope. Where *X* is the concentration, *Y* is the % response, top and bottom are plateaus in the same units as *Y*, EC_50_ is the 50% effective concentration, and hill slope is the slope factor.

### Cell Culture

22Rv1, DU145, and LNCaP cells were purchased from ATCC (22Rv1: CRL-2505; DU145: HTB-81) or from the UC Berkeley Cell Culture Facility, AR(v7)-HiBiT knock-in 22Rv1 cells were purchased from Promega Corp (CS3023257) and AIZ-AR stable 22Rv1 luciferase reporter cells were purchased from ABM (T3104). All three cell lines were cultured in RPMI-1640 medium containing 10% (v/v) fetal bovine serum (FBS) and 1× GlutMAX with or without 1X penicillin/streptomycin (Gibco). HEK293T cells were obtained from the UC Berkeley Cell Culture Facility and cultured in DMEM medium (containing 10% FBS and 1× GlutMAX), respectively. All the cell lines were maintained at 37°C with 5% CO_2_. Unless otherwise specified, all cell culture materials were purchased from Gibco. It is not known whether HEK293T cells are from male or female origin.

### Bortezomib and MLN4924 Rescue Studies

Cells were seeded in white 96-well plates (20,000 cells, 100 μL) and left overnight to adhere. The cells were pre-treated 24 hours after seeding with bortezomib (BTZ) or MLN4924 (1 μM or 0.2 μM final concentration, respectively) for 1 hour before treating with DMSO (vehicle) or a covalently acting small molecule. At 24 hours after compound treatment, reporter activity was assessed using the Nano-Glo® HiBiT lytic detection system (Promega, N3030). Cells were seeded in parallel for viability measurement using the CellTiter-Glo® 2.0 Cell Viability Assay (Promega, G9241) to normalize for differences in cell proliferation between conditions. Luminescence was detected using a Tecan Spark multimode microplate reader. Cells were seeded in 6-well plates (4 million cells, 2mL) and left overnight to adhere. The cells were pre-treated 24 hours after seeding with BTZ or MLN4924 (1 μM or 0.2 μM final concentration, respectively) for 1 hour before treating with DMSO (vehicle) or a covalently acting small molecule. At 24 hours after compound treatment, cells were harvested on ice and protein levels were assessed using Western blotting.

### Preparation of Cell Lysates

Cells were washed with cold PBS, scraped, and pelleted by centrifugation (1400*g*, 4 min, 4 °C). The lysates were then frozen at −80 °C or immediately processed for Western blotting. Pellets were resuspended in PBS (supplemented with Pierce protease inhibitor mini tablets, EDTA-free, Thermo Fisher Scientific, A32955) and sonicated on ice. Unless otherwise stated, total lysate was utilized for Western blotting and chemoproteomics experiments. When required, cells were clarified by centrifugation (20,000*g*, 10 min, 4 °C), and the lysate was transferred to new low-adhesion microcentrifuge tubes. Proteome concentrations were determined using BCA assay (Pierce, 23225), and the lysate was diluted to appropriate working concentrations.

### Western Blotting

For **Figures 1d, 1f, 1g, 1i, 2d, 2f, 5a, 5b, Figures S1a, S1c,** Western blotting experiments were done as follows. Antibodies to GAPDH (Proteintech, 60004-1-1g mouse) and AR (Cell Signaling Technology, D6F11 rabbit) were obtained and dilutions were prepared in accordance with the recommended manufacturer’s protocol. Proteins were resolved by sodium dodecyl sulfate poly(acrylamide) gel electrophoresis (SDS/PAGE) and transferred to nitrocellulose membranes using the Trans-Blot Turbo transfer system (Bio-Rad Laboratories, Inc.). Blots were blocked with 5% bovine serum albumin (BSA) in Tris-buffered saline containing Tween 20 (TBST) solution for 1 hour at room temperature, washed in TBST and probed with primary antibody diluted in diluent, as recommended by the manufacturer, overnight at 4 °C. Following washes with TBST, the blots were incubated in the dark with IR680- or IR800-conjugated secondary antibodies purchased from LI- CORbio and used at 1:10,000 dilution in 5% BSA in TBST at room temperature. Blots were visualized using an Odyssey Li-Cor fluorescent scanner after additional washes with TBST. If additional primary antibody incubations were required, the membrane was stripped using the ReBlot Plus strong antibody stripping solution (EMD Millipore, 2504), washed, and blocked again before being re-incubated with primary antibody. Blots were quantified and normalized to loading controls using ImageJ.

For **Figures 3d, 3f, 6a, Figure S2a,** Western blotting experiments were done as follows. Cells (5 x 10^5^) were plated in 6-well tissue-culture treated plates (Corning 3516) and returned to the incubator to adhere overnight. The following day, compounds or DMSO control were added to the cells and plates were returned to the incubator for 24 hours. Following, media was removed, cells were washed in 1X D-PBS (Gibco) and lysed by sonication in radioimmunoprecipitation assay (RIPA) buffer containing 1X NuPAGE LDS sample buffer (Thermo Fisher), complete protease inhibitor cocktail (Roche), and 1X NuPAGE sample reducing agent (Thermo Fisher). Samples were heat denatured at 70 °C for 10 minutes and then separated by SDS-PAGE on 4-20% Stain-free gradient gels (BioRad). Proteins were transferred to nitrocellulose blots using a Trans-Blot Turbo Transfer System (BioRad) and the blots were blocked for 30 minutes at room temperature in 1X Blocker FL Fluorescent Blocking Buffer (Thermo Fisher). Blots were then incubated overnight at 4 °C in primary antibodies diluted in 1X phosphate-buffered saline containing 0.02% Tween-20 (1X PBS-T) and 5% non-fat dry milk powder with rotation. The following day, blots were washed 3X in 1X PBS-T and then incubated in secondary antibodies diluted in 1X PBS-T plus 5% non-fat dry milk powder for 2 hours at room temperature. The blots were again washed 3X in 1X PBS-T and imaged on a ChemiDoc Imaging System (BioRad). Antibodies and dilutions used were as follows: rabbit anti-AR (1:1000, Cell Signaling Technology #5153), mouse anti-GAPDH (1:2000, UBP Bio #Y1041), IRDye 680RD goat anti-mouse secondary (1:10,000, LI-COR #926-68070), IRDye 800CW goat anti-rabbit secondary (1:10,000, LI-COR #926-32211).

### Purification of Recombinant AR-V7 Protein

Expression and purification of recombinant AR-v7 protein was performed by Selvita Life Sciences Solutions. A DNA sequence encoding amino acids 1-644 of the human androgen receptor was cloned into a plasmid vector with an N-terminal tobacco etch virus (TEV)-cleavable 6X-histidine (6His) tag. The plasmid was transformed into BL21(DE3) E. coli and cultured in TB medium supplemented with 1% glucose at 37 °C with 200 RPM rotation. The culture was grown to OD600 = 1, and protein expression was induced by the addition of isopropyl β-D-thiogalactopyranoside (IPTG) to 0.5 mM for 16 hours. Bacterial pellet was collected by centrifugation, resuspended in Lysis Buffer (50 mM HEPES pH 7.4, 300 mM NaCl, 1 mM TCEP, 5% glycerol, DNase I (10 µg/ml), 1X Protease Inhibitor Cocktail) and lysed by sonication. Lysate was subjected to centrifugation to pellet inclusion bodies containing insoluble protein. Inclusion bodies were washed by resuspension and centrifugation 3X in Wash Buffer I (50 mM HEPES pH 7.4, 300 mM NaCl, 1 mM TCEP, 1 % Triton X-100), followed by 3X washes in Wash Buffer II (50 mM HEPES pH 7.4, 300 mM NaCl, 1 mM TCEP). Inclusion bodies were subsequently solubilized in Denaturing Buffer (50 mM HEPES pH 7.4, 300 mM NaCl, 1 mM TCEP, 8 M Urea) and 6His-tagged protein was captured by affinity chromatography using Protino Ni2+- NTA agarose resin (Qiagen) under denaturing conditions. The resin was washed in AC Wash Buffer (50 mM HEPES pH 7.4, 300 mM NaCl, 1 mM TCEP, 8 M Urea, 20 mM imidazole) and target protein was eluted in Elution Buffer (50 mM HEPES pH 7.4, 300 mM NaCl, 1 mM TCEP, 8 M Urea, 300 mM imidazole). Denatured proteins were refolded by dialysis in Refolding Buffer (50 mM HEPES pH 7.4, 300 mM NaCl, 1 mM TCEP, 0.7 M L-arginine, 5% glycerol, 20 mM imidazole) and further purified by size exclusion chromatography (SEC) on a HiLoad 26/600 Superdex200pg column (GE Healthcare) in Storage Buffer (20 mM HEPES pH 7.4, 150 mM NaCl, 1 mM TCEP, 20 mM imidazole, 0.15 M L-arginine). Protein was concentrated by centrifugation over a 10k MWCO ultrafiltration disc (Merck Millipore), aliquots were flash frozen in liquid nitrogen, and protein was stored at −80 °C for later use.

### Gel-Based ABPP with AR-V7 Pure Protein

Gel-based ABPP methods were performed as previously described ^23^. Pure recombinant human His6, AR isoform 3 (AR-V7-like protein) was provided by Amgen Inc. AR-V7 (0.1 μg, 2.0 mg/mL) was diluted into 25 μL of phosphate-buffered saline (PBS) and 0.5 μL of either DMSO (vehicle) or covalently acting small molecule to achieve the desired concentration. After 30 min at 37 °C, the samples were treated with tetramethylrhodamine-5-iodoacetamide dihydroiodide (IA-rhodamine) (Setareh Biotech, 6222, prepared in anhydrous DMSO) (1 μM final concentration) for 1 hour at room temperature in the dark. Incubations were quenched by diluting the incubation with 10 μL of Laemmli SDS sample loading buffer, reducing 4× (Thermo Fisher Scientific, J60015.AC) and heated at 95 °C for 5 min. The samples were separated on precast 4−20% Criterion TGX gels (Bio-Rad Laboratories, Inc.). Fluorescent imaging was performed on a ChemiDoc MP (Bio-Rad Laboratories, Inc.). Inhibition of target labelling was assessed by densitometry using ImageJ. The imaged gels were stained using the Pierce Silver Stain Kit (Thermo Scientific, 24612) following the manufacturer’s instructions.

### AR Transcriptional Reporter Activity Rescue Studies

Cells were seeded in white 96-well plates (30,000 cells, 100 μL) and left overnight to adhere. The cells were transfected 24 hours after seeding with FLAG-tagged WT AR (OriGene Technologies, Inc., RC215316), C125S AR and AR-Luc (Signosis, Inc. LR-2105) plasmids. For plasmid transfection, 10 μL of the following mixture was added to the culture medium: 2.5 μg of WT AR or C125S AR plasmids and 2.5 μg of AR-Luc plasmid, together with 10 μL of X-tremeGENE HP DNA transfection reagent (Sigma-Aldrich, XTGHP-RO) in 500 μL of Opti-MEM. After 24 hours, cells were then treated with 10 nM 5α-dihydrotestosterone (DHT) solution (Sigma-Aldrich, D-073) and either DMSO (vehicle) or EN441. At 24 hours after compound treatment, reporter activity was assessed using the Bright-Glo® luciferase assay system (Promega, E2610). Cells were seeded in parallel for viability measurement using the CellTiter-Glo® 2.0 Cell Viability Assay (Promega, G9241) to normalize for differences in cell proliferation between conditions. Luminescence was detected using a Tecan Spark multimode microplate reader.

### AR Transcriptional Reporter Assay in 22Rv1 Cells

Androgen receptor transcriptional reporter assays were performed in white, opaque 96-well plates (Corning 3917) in 100 µL culture medium and [DMSO] = 0.1%. AIZ-AR stable 22Rv1 luciferase reporter cells (22Rv1 ARE-Luc, Applied Biological Materials) were dissociated in TrypLE (Gibco), cell number was determined using a Countess 3 Cell Counter (Thermo Fisher), and 2e4 cells per well were plated and returned to the incubator overnight. The following day, compounds or DMSO vehicle control were dispensed using a Multidrop Pico8 Digital Dispenser (Thermo Fisher) in biological triplicate and plates were returned to the incubator for 24 hours. Following, 100 µL prepared Steady-Glo luminescent reagent (Promega) was added to each well, plates were incubated for 5 minutes at room temperature with 300 RPM rotation and then equilibrated for an additional 3 minutes without rotation prior to reading. Luminescence was determined on a Perkin-Elmer Envision plate reader (Revvity). Values were normalized to DMSO control in Prism 10 (GraphPad) and curves were fit using a non-linear regression.

### AR-V7 HiBiT Cellular Assay

To assay the covalent ligand library, AR(v7)-HiBiT KI 22Rv1 were seeded at equal densities (20,000 cells) in white, opaque 96-well plates (Corning, 3917). The cells were treated 24 hours after seeding with either DMSO (vehicle) or a covalently acting small molecule. At 24 hours after compound treatment, reporter activity was assessed using the Nano-Glo® HiBiT lytic detection system (Promega, N3030) or the Steady-Glo luminescent reagent (Promega). Cells were seeded in parallel for viability measurement using the CellTiter-Glo® 2.0 Cell Viability Assay (Promega, G9241) to normalize for differences in cell proliferation between conditions. Luminescence was detected using a Tecan Spark multimode microplate.

### LC-MS/MS Analysis of EN1441 Reactivity with Pure AR-V7

Purified human His6, AR isoform 3 (AR-V7-like protein) (10 μg) in 50 μL PBS was incubated with EN1441 (50 μM) for 1 hour at 37 °C. The sample was precipitated by addition of 10 μL of 100% (w/v) trichloroacetic acid, before cooling to −80 °C overnight. The sample was then spun at max speed for 10 min at 4 °C, supernatant was carefully removed, and the sample was washed three times with 200 μL of ice-cold 0.01 M HCl/90% acetone solution, with spinning at max speed for 5 min at 4 °C between washes. The pellet was resuspended in 30 μL of 8 M urea in PBS and 30 μL of ProteaseMax surfactant (20 μg/mL in 100 mM ammonium bicarbonate, Promega Corp, V2071) with 15 sec vortexing. Ammonium bicarbonate (40 μL, 100 mM) was then added for a final volume of 100 μL. The sample was reduced with 10 μL of 110 mM TCEP (10 mM final concentration) at 60 °C for 30 min and alkylated with 10 μL of 150 mM iodoacetamide (12.5 mM final concentration) at 37 °C for 30 min. The sample was then diluted with 120 μL of PBS before 1.2 μL of ProteaseMax surfactant (0.1 mg/mL in 100 mM ammonium bicarbonate) and sequencing grade trypsin (1 μL, 0.5 mg/mL in 50 mM ammonium bicarbonate, Promega Corp, V5111) were added for overnight incubation at 37 °C. The next day, the sample was acidified with 12 μL formic acid, spun at 13,200 g at 4 °C for 30 min, and the supernatant was transferred to a new tube and stored at −80 °C until mass spectrometry analysis. Mass spectrometry samples were prepped by fractionating using high pH reversed-phase peptide fractionation kits (Thermo Fisher Scientific, 84868) according to manufacturer’s protocol.

### Quantitative Tandem Mass Tagging (TMT)-Based Proteomic Profiling of EN1441

22Rv1 cells were treated when they reached 80% confluency, with either DMSO (vehicle) or 50 μM of EN1441 for 16 hours. Total cell lysates were prepared using sonication, and proteome concentration was normalized to 100 μg of protein per replicate in 100 μL with PBS. To prevent preliminary precipitation, 0.5 μL of 10% sodium dodecyl sulfate (SDS) was added to each sample. Each sample was reduced with 5 μl of 200 mM TCEP (55 °C, 1 hour) and alkylated with 5 μl of 375 mM iodoacetamide (25 °C, 30 min in darkness). After the reaction, the protein samples were precipitated overnight in acetone at −20 °C. After the overnight precipitation, the samples were centrifuged at 8,000 g for 10 minutes at 4 °C. The supernatant was removed, and the precipitated protein samples were resuspended in 100 μL of 50 mM tetraethylammonium bromide (TEAB) buffer. 2.5 μL of 1.0 mg/mL sequencing grade trypsin (Promega, V5111) and 1 μL of 100 mM calcium chloride in H_2_O were then added, and the proteins were digested overnight at 37 °C with shaking. Peptide amounts were quantified using a quantitative colorimetric peptide assay (Thermo Fisher Scientific Pierce, 23275) and the individual normalized 100 μL samples were then labelled with isobaric tags using a commercially available Tandem Mass Tag™ (TMT) 6-plex kit (TMTsixplex™) (Thermo Fisher Scientific, 90061), in accordance with the manufacturer’s protocol. TMT samples were then consolidated and separated using high pH reversed-phase peptide fractionation kits (Thermo Fisher Scientific, 84868) according to the manufacturer’s protocol. Fractions were vacuum concentrated, then reconstituted in 0.1% (v/v) formic acid and centrifuged at 20,000 g (5 min) in preparation for LC-MS/MS analysis.

### Mass Spectrometry Analysis

Mass spectrometry analysis was performed on an Orbitrap Eclipse Tribrid Mass Spectrometer with a High Field Asymmetric Waveform Ion Mobility (FAIMS Pro) Interface (Thermo Fisher Scientific) with an UltiMate 3000 Nano Flow Rapid Separation LCnano System (Thermo Fisher Scientific). Offline fractionated samples (5 μL aliquot of 25 μL sample) were injected via an autosampler (Thermo Fisher Scientific) onto a 5 μL sample loop, which was subsequently eluted onto an Acclaim PepMap 100 C18 HPLC column (75 μm × 50 cm, NanoViper). The peptides were separated at a flow rate of 0.3 μL/min using the following gradient: 2% buffer B (100% acetonitrile with 0.1% formic acid) in buffer A (95:5 water/acetonitrile, 0.1% formic acid) for 5 min, followed by a gradient from 2−40% buffer B from 5−159 min, 40−95% buffer B from 159−160 min, held at 95% B from 160−179 min, 95% to 2% buffer B from 179−180 min, and then 2% buffer B from 180−200 min. The voltage applied to the nano-LC electrospray ionization source was 2.1 kV. Data were acquired through an MS1 master scan (Orbitrap analysis, resolution 120,000, 400−1800 *m*/*z*, RF lens 30%, heated capillary temperature 250 °C) with dynamic exclusion (repeat count 1, duration 60 sec). Data-dependent data acquisition comprised a full MS1 scan, followed by sequential MS2 scans based on 2 sec cycle times. FAIMS compensation voltages (CVs) of −35, −45, and −55 were applied. MS2 analysis consisted of a quadrupole isolation window of 0.7 *m*/*z* of the precursor ion followed by a higher energy collision dissociation (HCD) energy of 38% with an orbitrap resolution of 50,000.

For quantitative TMT-based proteomic analysis, data was extracted from raw files in the form of MS1, MS2 and MS3 files using RAW Xtractor (version 1.9.9.2; publicly available at http://fields.scripps.edu/downloads.php) and searched against the Uniprot human database using ProLuCID search methodology in IP2 v.3−v.5 (Integrated Proteomics Applications, Inc.) ^53^. Trypsin cleavage specificity (cleavage at K, R, except if followed by P) allowed for up to 2 missed cleavages. ProLucid searches allowed for static modification of cysteine residues (+57.02146 due to alkylation), methionine oxidation (+15.9949) and TMT modification of N-terminus, and lysine residues were set as variable modifications. Reporter ion ratio calculations were performed using summed abundances with the most confident centroid selected from the 20 ppm window. Only peptide-to-spectrum matches that are unique to a given identified protein within the total data set are considered for protein quantitation. High-confidence protein identifications were reported with a <1% false discovery rate (FDR) cut-off. Differential abundance significance was estimated using ANOVA with Benjamini−Hochberg correction to determine the *p*-values.

### *In situ* Alkyne Probe Labelling and Endogenous AR/AR-V7 Pulldown

22Rv1 cells were pre-treated 24 hours after seeding with bortezomib (BTZ) (1 μM final concentration) for 1 hour before treating with either DMSO (vehicle) or 5 μM EN1441-alkyne probes for 4.5 hours. Cells were collected in PBS (supplemented with Pierce protease inhibitor mini tablet) and lysed by sonication. For preparation of Wester blotting samples, the lysate (2.5 mg protein in 1 mL) was aliquoted into low retention microcentrifuge tubes and the following were then added: 10 μl of 10 mM biotin picolyl azide (Sigma-Aldrich, 900912), 10 μl of TCEP (14.0 mg/mL in H_2_O), 30 μl of TBTA (0.9 mg/mL in 1:4 in DMSO/*t*-BuOH) and 10 μl of CuSO_4_ (12.5 mg/mL in H_2_O). Samples were incubated for 1 hour at room temperature. Following CuAAC, proteins were precipitated by centrifugation at 6,500 g and washed twice with ice-cold methanol (500 μL). The samples were spun in a prechilled (4 °C) centrifuge at 6,500 g for 4 min, and the methanol supernatant was removed. The protein pellet was subsequently reconstituted in 200 μL PBS containing 1.2% sodium dodecyl sulfate (SDS) (w/v) by probe sonication. The resultant solution was heated to 90 °C for 5 min (10 μL was saved for input) and diluted with 0.5 mL PBS. To each sample, 70 μL of high-capacity streptavidin agarose resin (50% slurry, Thermo Fisher Scientific, 20359) was added using 2 × 250 μL PBS and incubated overnight on a rocker at four °C. The bead mixtures were spun at 1400 g for 3 min, and the supernatant was removed. The beads were then transferred to spin columns and washed with 1 mL of 0.2% SDS (w/v), 3 × 1 mL PBS, and 3 × 1 mL H_2_O. Proteins were eluted by boiling the beads in PBS and 20 μL of Laemmli SDS sample loading buffer, reducing 4× for 15 min at 95 °C.

For TMT-based quantitative proteomic profiling, the washed beads were suspended in 500 μL of 6M urea and incubated with 25 μL of dithiothreitol (DTT, Invitrogen, 15508-013) (30 mg/mL) at 65 °C for 20 min. 25 μL of 400 mM iodoacetamide (GBiosciences, #0461) (74 mg/mL) were then added and the samples were incubated at 37 °C for 30 min. Centrifugation of the bead mixture at 1400 g for 2 min was followed by the aspiration of the supernatant. The beads were then resuspended in 100 of 50 mM tetraethylammonium bromide (TEAB) buffer to which 4 μL of 0.5 mg/mL sequencing grade trypsin (Promega, V5111A) and 1 μL of 100 mM calcium chloride in H_2_O were added, and the bead bound proteins were digested overnight at 37 °C with shaking.

Peptide amounts were quantified using a quantitative colorimetric peptide assay (Thermo Fisher Scientific Pierce, 23275) and the individual samples were then labelled with isobaric tags using a commercially available Tandem Mass Tag™ (TMT) 6-plex kit (TMTsixplex™) (Thermo Fisher Scientific, 90061), in accordance with the manufacturer’s protocol. TMT samples were then consolidated and separated using high pH reversed-phase peptide fractionation kits (Thermo Fisher Scientific, 84868) according to the manufacturer’s protocol. Fractions were vacuum concentrated, then reconstituted in 0.1% (v/v) formic acid for LC-MS/MS analysis.

### Thermal Denaturation Sensitivity Assay

22Rv1 cells were seeded in 6 cm^2^ plates at 2,500,000 cells per well (4 mL total) and left overnight to adhere. The cells were then treated with DMSO (vehicle) or 50 μM EN1441 for 1 hour. The cells were harvested, washed with PBS, then suspended in 1 mL of PBS (supplemented with Pierce protease inhibitor mini tablet) and maintained with DMSO or 50 μM EN1441. Each sample set was divided equally amongst the eight tubes of a PCR strip (i.e., 100 μL volume per tube), and each tube was designated a temperature (45.0 °C, 46.5 °C, 49.2 °C, 53.3 °C, 58.2 °C, 62.4 °C, 65.2 °C, 66.9 °C). Samples were heated at their respective temperatures for 3 min in a 96-well thermal cycler (Bio-Rad, T100), then incubated at 25 °C for another 3 min. Cells were then immediately snap-lysed in liquid nitrogen (three freeze-thaw cycles). Cell debris along with precipitated and aggregated proteins were removed by centrifugation at 20,000 g for 20 min at 4 °C. 80 μL of the supernatant was transferred into a new tube and following the addition of 16 μL Laemmli SDS sample loading buffer, reducing 4× was heated to 95 °C for 5 min. Samples were analyzed by Western blotting analysis.

For the thermal denaturation sensitivity assay analysis of EN1441 on pure AR-V7 protein, human His6, AR isoform 3 (AR-V7-like protein) (20 μg) in 1 mL PBS was incubated with DMSO (vehicle) or EN1441 (50 μM) for 1 hour at 37 °C. This represents a single sample set. The experiment followed that already previously described above using the following temperatures: 55.1 °C, 56.8 °C, 59.6 °C, 63.6 °C, 68.6 °C, 73.4 °C, 76.3 °C and 78.0 °C.

For thermal denaturation sensitivity assay based-TMT analysis, 22Rv1 cells were seeded in 15 cm^2^ plates and left overnight to adhere. The cells (∼80% confluency) were then treated with DMSO (vehicle) or 50 μM EN1441 for 1 hour. The cells were harvested, washed with PBS, then suspended in PBS (supplemented with Pierce protease inhibitor mini tablet). Cellular lysates were obtained by sonication, then cleared by centrifugation at 16,000 g for 10 min at 4 °C, and the proteome concentration was normalized to 1.3 mg/mL. Each replicate (1 mL) was divided into equal portions for temperature treatment (45.0 °C, 48.8 °C, 57.1 °C, 63.4 °C, 65.0 °C, 69.0 °C, 76.7 °C, 83.6 °C), which was performed for 3 min at the desired temperature in a 96- well thermal cycler. The samples were then incubated at room temperature for 6 min before combining each temperature point for every replicate. Cells were clarified by centrifugation (20,000*g*, 45 min, 4 °C), and the lysate was transferred to new low-adhesion microcentrifuge tubes. Proteome concentrations were determined using BCA assay (Pierce, 23225) and normalised across all six sample sets. 200 μg of protein from each replicate was reduced with TCEP (10 μl of 200 mM), alkylated with iodoacetamide (10 μl of 375 mM) and tryptically digested (5 μg of trypsin) overnight as previously described. The individual samples were then labelled with a commercially available Tandem Mass Tag™ (TMT) 6-plex kit (TMTsixplex™) (Thermo Fisher Scientific, 90061) in accordance with the manufacturer’s protocol. Downstream proteomics sample processing was performed as previously described in the TMT proteomics section.

### Site-Directed Mutagenesis on FLAG-Tagged Wild-Type AR Plasmid

FLAG-tagged wild type AR plasmid was purchased from OriGene Technologies, Inc. (RC215316), and site-directed mutagenesis was performed using the Q5-site-directed mutagenesis kit (New England biolabs, E0554S) in accordance with the manufacturer’s protocol. The following primer pair sequences were designed using the New England Base Changer tool:

C125S Forward Primer: GGC CCT GGA GAG CCA CCC CGA GA
C125S Reverse Primer: GAC TGC GGC TGT GAA GGT TGC TG

The mutant plasmid was transformed into 5-alpha Competent *E*. *coli* cells (New England biolabs, C2987H) and the cells grown on LB agar Kanamycin plates (Sigma Aldrich L0543). Colonies selected were then cultured in LB broth (Thermo Fisher Scientific, BP1426-500) with 100 μg/mL Kanamycin (100 mg/mL) overnight at 200 rpm at 37 °C. *E*. *coli* containing the desired plasmids were pelleted, lysed, and neutralized using the commercially available Monarch^®^ plasmid DNA miniprep kit (New England biolabs, T1010) in accordance with the manufacturer’s protocol. The eluted plasmid concentration was determined using Nanodrop quantification.

### Primers for Mutant Plasmid Sequence Confirmation

Plasmids were sent to ELIM Biopharmaceuticals, Inc. to confirm mutant sequences. The following primer was used:

T7 Forward: TAATACGACTCACTATAGGG

### Expression of FLAG-Tagged Wild-Type AR and FLAG-Tagged C125S Mutant

For rescue studies, HEK293T cells were seeded in 6-well plates at 1,500,000 cells per well (2 mL total). The cells were transfected 24 hours after seeding with FLAG-tagged WT AR (OriGene Technologies, Inc., RC215316) and C125S AR plasmids. For plasmid transfection, 1.25 μg of overexpression plasmid was added to the culture medium together with 3.75 μL of the Lipofectamine™ 3000 transfection reagent and 5 μL P3000 (Invitrogen, L3000001) in 250 μL of reduced serum medium, Opti-MEM (Thermo Fisher Scientific, 31985062). After another 24 hours, the cells were treated with DMSO (vehicle) or 50 μM EN1441 for 6 hours. The cells were harvested, washed with PBS, then suspended in 1 mL of PBS (supplemented with Pierce protease inhibitor mini tablet). Total cell lysates were prepared using sonication and protein levels were assessed using Western blotting.

### Immunofluorescence and Microscopy

Immunofluorescence and Microscopy. For stress granule assessment, 22Rv1 cells were seeded in 6-well plates at 2,000,000 cells per well (2 mL total) and left to adhere overnight. The cells were then transfected with pEGFP-G3BP1-WT plasmid (Addgene, #135997). For plasmid transfection, 0.75 μg of overexpression plasmid was added to each well together the Lipofectamine™ 3000 transfection reagent and P3000 (Invitrogen, L3000001) in accordance with the manufacturer’s protocol. After 24 hours, the cells were re-seeded in a 4- compartment cell culture dish (Greiner Bio-One, 627870). After another 24 hours, the cells were then treated with DMSO (vehicle) or EN1441 for 4 hours. The cells were then fixed with 4% paraformaldehyde, methanol free (Cell Signaling, #47746) for 15 min, washed with PBS three times, permeabilized with 0.25% tris buffered saline with Tween20 (TBST) for 15 min, and then stained with 2 mM Hoechst stain (Invitrogen, H3570), with all steps performed at room temperature. Images were captured using a Zeiss LSM880 FCS microscope with a 40× oil objective.

### Viability assays

Viability assays were performed in white, opaque 96-well plates (Corning 3917) in 100 µL culture medium and [DMSO] = 0.1%. 1e4 cells were plated per well and returned to the incubator overnight to adhere. The following day, compounds or DMSO vehicle control were dispensed using a Multidrop Pico8 Digital Dispenser (Thermo Fisher) in biological triplicate and plates were returned to the incubator for 72 hours. Following, 100 µL prepared Cell Titer Glo luminescent reagent (Promega) was added to each well, plates were incubated for 5 minutes at room temperature with 300 RPM rotation and then equilibrated for an additional 3 minutes without rotation prior to reading. Luminescence was determined on a Perkin-Elmer Envision plate reader (Revvity). Values were normalized to DMSO control in Prism 10 (GraphPad) and curves were fit using a non-linear regression.

### Detergent Solubility Assays

5e5 cells were plated in 6-well tissue culture-treated plates (Corning 3516) and returned to the incubator to adhere overnight. The following day, compounds or DMSO control were added to the cells and plates were returned to the incubator for 24 hours. Following, media was removed, cells were washed in 1X D-PBS (Gibco) and lysed by sonication in ice-cold PBS containing 3% 3-((3-cholamidopropyl) dimethylammonio)-1- propanesulfonate (CHAPS, Thermo Fisher) and cOmplete protease inhibitor cocktail. A sample representing the “total” was immediately collected and prepared for western blotting as described previously. The remaining lysate was then centrifuged for 30 minutes at 13000 RPM at 4 °C, and the supernatant was collected and prepared for western blotting representing the “soluble” component. The pellet was washed in ice-cold lysis buffer, and again centrifuged for 10 minutes at 13000 RPM at 4 °C. The wash buffer was removed, and the pellet representing the “insoluble” component was resuspended in lysis buffer containing 1X NuPAGE LDS sample buffer, cOmplete protease inhibitor cocktail, and 1X NuPAGE sample reducing agent and sonicated to disrupt LDS-insoluble aggregates. All samples were heat denatured at 70 °C for 10 minutes and analyzed by western blot as described previously.

### RT-qPCR

3e5 cells were plated in 12-well tissue culture-treated plates (Corning 3513) and returned to the incubator to adhere overnight. The following day, compounds or DMSO control were added to the cells in biological triplicate and plates were returned to the incubator for 20 hours. Following, media was removed, cells were washed in 1X D-PBS (Gibco), and RNA was collected using the RNeasy Plus RNA purification kit (Qiagen) according to the manufacturer’s instructions. RNA concentrations were determined using a NanoDrop Spectrophotometer (Thermo Fisher), and 1 µg of RNA was converted to cDNA using the Quantitect Reverse Transcription Kit (Qiagen) according to the manufacturer’s instructions. qPCR reactions were prepared in technical triplicate in MicroAmp 384-well qPCR plates (Applied Biosystems) with 10 µL final volume using PrimeTime 2X Gene Expression Master Mix (Integrated DNA Technologies). *KLK3* and *ACTB* transcript levels were detected using predesigned PrimeTime qPCR Probe Assays (Integrated DNA Technologies). Thermal cycling was performed on a QuantStudio 12K Flex Real-Time PCR System (Thermo Fisher) using standard cycling conditions. Relative fold-change was determined using the 2^-ddC_t_ method and normalized to DMSO control.

### RNAseq Library Preparation

Total RNA (500ng) was used to generate cDNA libraries using the QuantSeq 3’ mRNA-Seq V2 Library Prep Kit (Lexogen, catalog #191.96) according to the manufacturer’s instructions. The workflow included cDNA synthesis, library amplification with 15 PCR cycles, and final bead-based purification. The resulting cDNA libraries were quantified using Qubit and Agilent 4200 TapeStation. Final sequencing libraries were pooled at equimolar concentrations and sequenced as 150 bp paired end reads on the NovaSeq 6000 platform (Illumina).

### Data analysis

For the 3’ mRNA-seq data, each library produced over 50 million reads, with only the Read 1 FASTQ files used for downstream analysis. Quality control, trimming, alignment, and other preprocessing steps were carried out using the nf-core/RNAseq pipeline. After quality control, reads were aligned to the human reference genome (hg38) with the STAR aligner (v2.7.10), and gene-level read counts were quantified using Salmon (v1.10.1). Raw read counts > 100 in more than three samples were selected for downstream analysis. Differential gene expression analysis was conducted using DESeq2 (1.42.1) and a threshold of |log2(FoldChange)| > 1 and an adjusted p-value < 0.05 was applied to identify differentially expressed genes.

