## Supporting Information for "Covalent Destabilizing Degrader of AR and AR-V7 in Androgen-Independent Prostate Cancer Cells"

**Supporting Table Legends**

**Table S1. Cellular covalent ligand screening to discover AR-V7 degraders.** Covalent ligand screen in 22Rv1 cells expressing an endogenously C-terminally tagged HiBiT peptide on AR-V7. These cells were treated with DMSO vehicle or cysteine-reactive covalent ligand (50 μM) for 24 h, after which HiBiT-AR-V7 was detected by luminescence through detection with LgBiT. Structures of compounds screened are in Tab 1. Values for individual compounds are noted as the treatment/control ratio and are shown in Tab 2. Screening data is from n=1 per compound.

**Table S2. Quantitative tandem mass tagging (TMT)-based proteomic profiling of EN1441.** 22Rv1 cells were treated with DMSO vehicle or EN1441 (50 μM) for 16 h, and resulting cell lysates were subjected to quantitative proteomic profiling. Data are from n=3 biologically independent replicates per group.

**Table S3. EN1441-alkyne pulldown chemoproteomic profiling.** 22Rv1 cells were pre-treated with BTZ (1 μM) for 1 h prior to treatment of cells with DMSO vehicle or EN1441-alkyne (5 μM, 4.5 h). Resulting lysates were subjected to CuAAC with an azide-functionalized biotin handle, and probe-modified proteins were avidin-enriched, eluted, and proteins were analyzed by TMT-based quantitative proteomic profiling. Data are from n=3 biologically independent replicates per group.

**Table S4. Transcriptomic profiling of EN1441.** Transcriptomic profiling of enzalutamide, EN1441, and CMZ139 in 22Rv1 cells. 22Rv1 cells were treated with 5 μM enzalutamide, EN1441, or CMZ139 for 20 h, after which extracted RNA was subjected to RNA sequencing and transcripts were quantified. Tab 1 shows normalized gene counts for RNAseq data from all treatment groups. Tabs 2-4 show fold-changes and statistical assessment for enzalutamide, EN1441, or CMZ139 versus DMSO vehicle treatment, respectively. Tab 5 shows treated/control individual replicate ratios for AR response genes. Data are from n=3 biologically independent replicates per group.

**Table S5.** CETSA-TMT proteomic profiling on EN1441. 22Rv1 cells were treated with DMSO vehicle or EN1441 (50 μM) for 1 h, after which cells were heated over a range of temperatures, insoluble proteins were removed, and remaining proteomes from control temperature groups were combined, treated temperature groups were combined, and subjected to TMT-based quantitative proteomics to identify proteins that were destabilized by EN1441 treatment. Data are from n=3 biologically independent replicates per group.

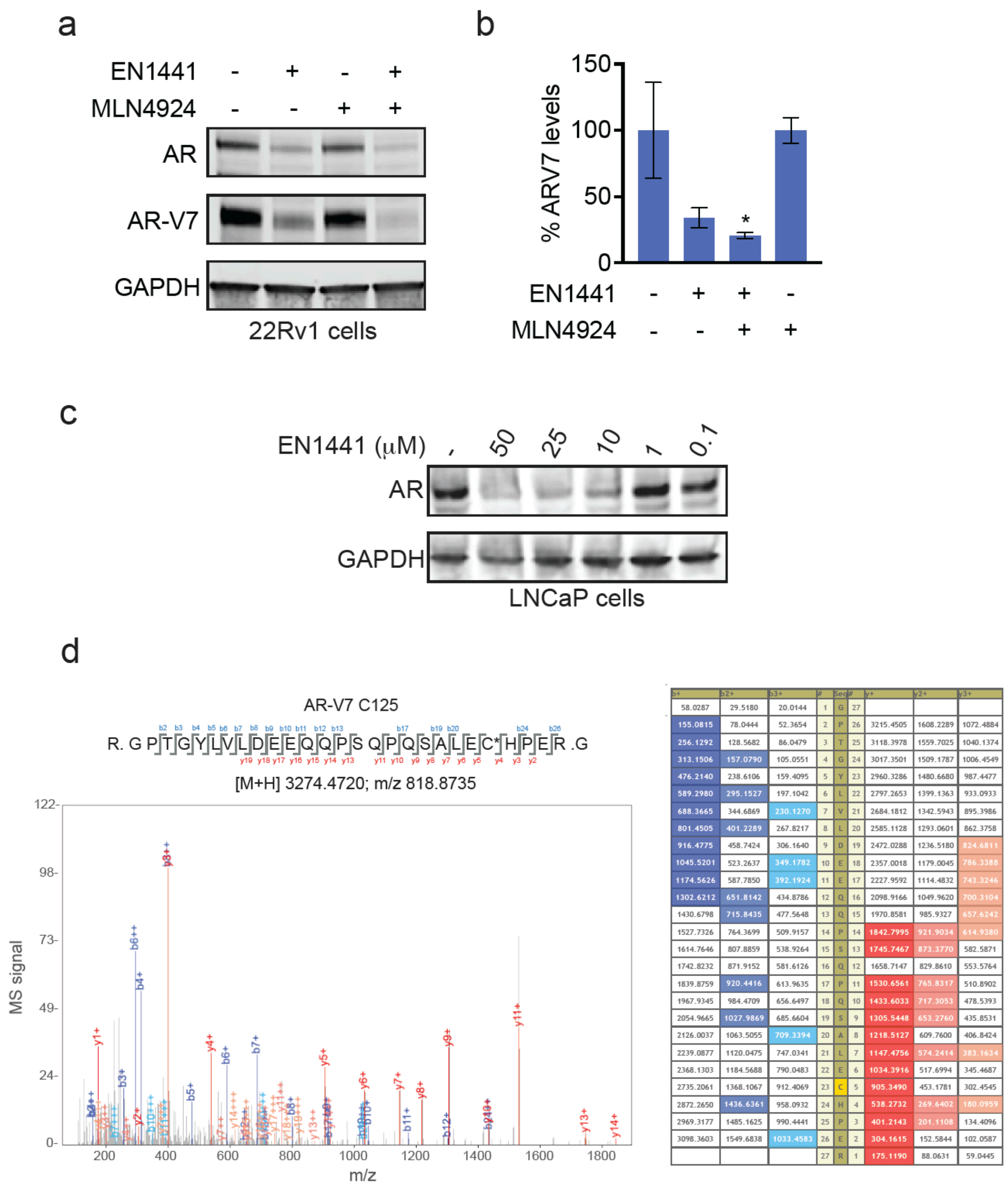


| b+ | b2+ | b3+ | # | Seq | # | y+ | y2+ | y3+ |
|---|---|---|---|---|---|---|---|---|
| 58.0287 | 29.5180 | 20.0144 | 1 | G | 27 | | | |
| 155.0815 | 78.0444 | 52.3654 | 2 | P | 26 | 3215.4505 | 1608.2289 | 1072.4884 |
| 256.1292 | 128.5682 | 86.0479 | 3 | T | 25 | 3118.3978 | 1559.7025 | 1040.1374 |
| 313.1506 | 157.0790 | 105.0551 | 4 | G | 24 | 3017.3501 | 1509.1787 | 1006.4549 |
| 476.2140 | 238.6106 | 159.4095 | 5 | Y | 23 | 2960.3286 | 1480.6680 | 987.4477 |
| 589.2980 | 295.1527 | 197.1042 | 6 | L | 22 | 2797.2653 | 1399.1363 | 933.0933 |
| 688.3665 | 344.6869 | 230.1270 | 7 | V | 21 | 2684.1812 | 1342.5943 | 895.3986 |
| 801.4505 | 401.2289 | 267.8217 | 8 | L | 20 | 2585.1128 | 1293.0601 | 862.3758 |
| 916.4775 | 458.7424 | 306.1640 | 9 | D | 19 | 2472.0288 | 1236.5180 | 824.6811 |
| 1045.5201 | 523.2637 | 349.1782 | 10 | E | 18 | 2357.0018 | 1179.0045 | 786.3388 |
| 1174.5626 | 587.7850 | 392.1924 | 11 | E | 17 | 2227.9592 | 1114.4832 | 743.3246 |
| 1302.6212 | 651.8142 | 434.8786 | 12 | Q | 16 | 2098.9166 | 1049.9620 | 700.3104 |
| 1430.6798 | 715.8435 | 477.5648 | 13 | Q | 15 | 1970.8581 | 985.9327 | 657.6242 |
| 1527.7326 | 764.3699 | 509.9157 | 14 | P | 14 | 1842.7995 | 921.9034 | 614.9380 |
| 1614.7646 | 807.8859 | 538.9264 | 15 | S | 13 | 1745.7467 | 873.3770 | 582.5871 |
| 1742.8232 | 871.9152 | 581.6126 | 16 | Q | 12 | 1658.7147 | 829.8610 | 553.5764 |
| 1839.8759 | 920.4416 | 613.9635 | 17 | P | 11 | 1530.6561 | 765.8317 | 510.8902 |
| 1967.9345 | 984.4709 | 656.6497 | 18 | Q | 10 | 1433.6033 | 717.3053 | 478.5393 |
| 2054.9665 | 1027.9869 | 685.6604 | 19 | S | 9 | 1305.5448 | 653.2760 | 435.8531 |
| 2126.0037 | 1063.5055 | 709.3394 | 20 | A | 8 | 1218.5127 | 609.7600 | 406.8424 |
| 2239.0877 | 1120.0475 | 747.0341 | 21 | L | 7 | 1147.4756 | 574.2414 | 383.1634 |
| 2368.1303 | 1184.5688 | 790.0483 | 22 | E | 6 | 1034.3916 | 517.6994 | 345.4687 |
| 2735.2061 | 1368.1067 | 912.4069 | 23 | C | 5 | 905.3490 | 453.1781 | 302.4545 |
| 2872.2650 | 1436.6361 | 958.0932 | 24 | H | 4 | 538.2732 | 269.6402 | 180.0959 |
| 2969.3177 | 1485.1625 | 990.4441 | 25 | P | 3 | 401.2143 | 201.1108 | 134.4096 |
| 3098.3603 | 1549.6838 | 1033.4583 | 26 | E | 2 | 304.1615 | 152.5844 | 102.0587 |
| | | | 27 | R | 1 | 175.1190 | 88.0631 | 59.0445 |

**Figure S1. Characterization of EN1441. (a)** AR and AR-V7 levels in 22Rv1 cells treated with EN1441. 22Rv1 parental cells were pre-treated with DMSO vehicle or MLN4924 (0.2 μM) for 1 h prior to treatment of cells with DMSO vehicle or EN1441 (50 μM) for 24 h, and AR and AR-V7 levels and loading control GAPDH were detected by Western blotting. **(b)** Quantification of experiment in **(a)**. **(c)** Dose-response of AR degradation with EN1441 treatment. LNCaP cells were treated with DMSO or EN1441 for 8 h, and AR and GAPDH levels were detected by Western blotting. **(d)** EN1441 modifies intrinsically disordered C125 on AR-V7. Pure AR-V7 protein was incubated with EN1441 (50 μM, 1 h) after which the reaction was quenched, the protein was tryptically digested and peptides were analyzed by LC-MS/MS and the EN1441-adducted tryptic peptide was identified. Shown is the MS/MS coverage of the EN1441-adducted AR-V7 peptide and the coverage of b- and y-ions. Data from **(a-d)** are from n=3 biologically independent replicates per group. Blots and spectra in **(a,c,d)** are representative. Bar graph in **(b)** shows average ± sem with significance expressed as *p<0.05 compared to vehicle-treated controls.

a

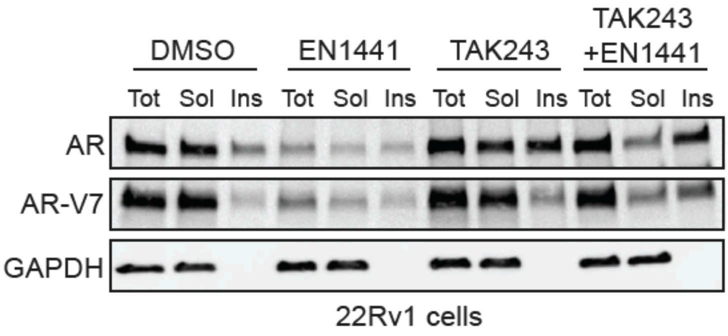


b

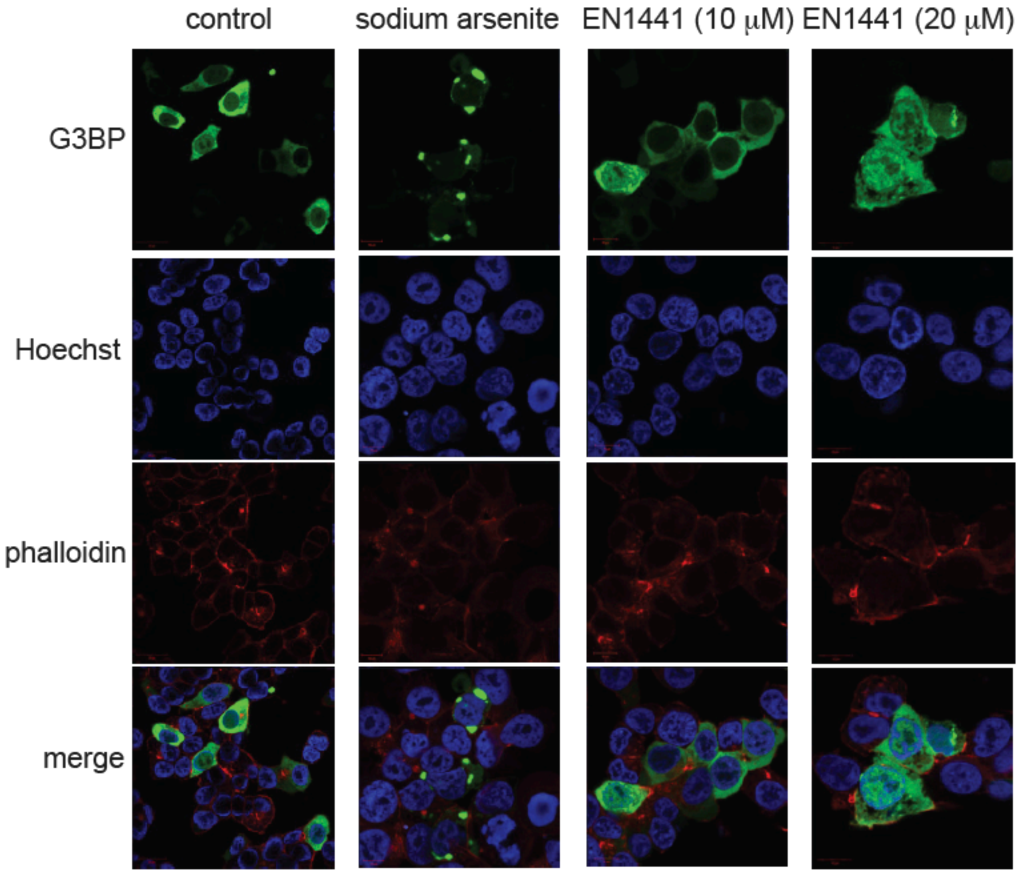


**Figure S2. EN1441 does not cause stress granules but does aggregate and degrade AR and AR-V7 in a ubiquitin-dependent manner. (a)** Total, soluble, and insoluble AR, AR-V7, and loading control GAPDH levels in 22Rv1 cells. 22Rv1 cells were pre-treated with DMSO or TAK243 (1 μM) for 1 h prior to treatment of cells with DMSO vehicle or EN1441 (20 μM) for 24 h. Soluble (sol) and insoluble (ins) fractions were isolated from total cell lysate (tot), separated on SDS/PAGE, and AR, AR-V7, and GAPDH levels were detected by Western blotting. **(b)** 22Rv1 cells were treated with DMSO vehicle control, stress granule-causing sodium arsenite (200 μM), or EN1441 for 4 h. Stress granules puncta were detected by staining for G3BP, nuclei were stained with Hoechst, and actin was stained with phalloidin.

### Synthetic Methods and Characterization

#### 6-chloro-1-hex-5-ynyl-3,3-dimethyl-4-prop-2-enoyl-quinoxalin-2-one (EN1441-Alkyne)

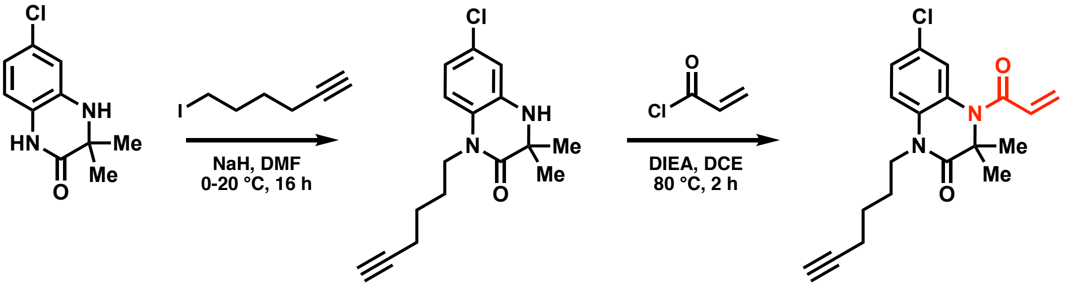


1 2 **EN1441-Alkyne**

*Step 1*: To a solution of 6-chloro-3,3-dimethyl-1,4-dihydroquinoxalin-2-one (200 mg, 949.40 μmol) in DMF (3 mL) was added NaH (45.57 mg, 1.14 mmol, 60% purity) at 0 °C under $N_2$. After stirring for 30 min at 0 °C, 6-iodohex-1-yne (237.01 mg, 1.14 mmol) was added. The reaction was then stirred at 20 °C for 16 h under $N_2$. The reaction mixture was then quenched by the addition of $H_2O$ (5 mL) at 0 °C, and then extracted with ethyl acetate (3 mL × 3). The combined organic layers were then washed with brine (5 mL), dried over $Na_2SO_4$, filtered, and concentrated *in vacuo*. Purification *via* prep-HPLC (column: Waters Xbridge BEH C18, 10 μm, 100 × 25 mm; mobile phase: [$H_2O$ (10 mM $NH_4HCO_3$) – acetonitrile]; gradient: 40% – 80% B over 8.0 min) gave 6-chloro-1-hex-5-ynyl-3,3-dimethyl-4H-quinoxalin-2-one (190 mg, 653.40 μmol, 68.8% yield) as a white solid. MS (ESI) *m/z* = 291.0 $[M+H]^+$.

*Step 2*: To a solution of 6-chloro-1-hex-5-ynyl-3,3-dimethyl-4H-quinoxalin-2-one (80 mg, 275.12 μmol) in 1,2-dichloroethane (3 mL) was added dropwise *N*,*N*-diisopropylethylamine (142.22 mg, 1.10 mmol, 191.68 μL) and prop-2-enoyl chloride (62.25 mg, 687.79 μmol, 55.88 μL) and the mixture was stirred at 80 °C for 2 h. The reaction mixture was then concentrated *in vacuo*. Purification *via* prep-HPLC (column: Welch Ultimate C18, 5 μm, 120 × 30 mm; mobile phase: [$H_2O$ (0.1% v/v TFA) – acetonitrile]; gradient: 60% – 90% B over 8.0 min) gave 6-chloro-1-hex-5-ynyl-3,3-dimethyl-4-prop-2-enoyl-quinoxalin-2-one (46.7 mg, 134.07 μmol, 48.7% yield, 99.0% purity) as a yellow solid. MS (ESI) *m/z* = 345.1 $[M+H]^+$; $\delta_H$ (400 MHz, DMSO-$d_6$) 7.34 (s, 2H), 7.03 (s, 1H), 6.31 – 6.17 (m, 2H), 5.75 (dd, *J* = 4.2, 7.6 Hz, 1H), 3.93 (t, *J* = 7.2 Hz, 2H), 2.77 (t, *J* = 2.6 Hz, 1H), 2.18 (dt, *J* = 2.4, 6.8 Hz, 2H), 1.65 (quin, *J* = 7.4 Hz, 2H), 1.50 (s, 6H), 1.48 – 1.39 (m, 2H). $\delta_C$ (101 MHz, DMSO-$d_6$) 169.55, 166.08, 132.15, 130.26, 128.86, 128.72, 126.48, 125.76, 124.23, 116.88, 84.17, 71.51, 60.78, 41.55, 25.41, 25.09, 22.92, 17.34. MS (ESI-TOF) Calcd for $C_{19}H_{22}ClN_2O_2^+$ $[M+H]^+$ : 345.1364; found: 345.1378.

**6-chloro-3,3-dimethyl-4-propanoyl-1H-quinoxalin-2-one**

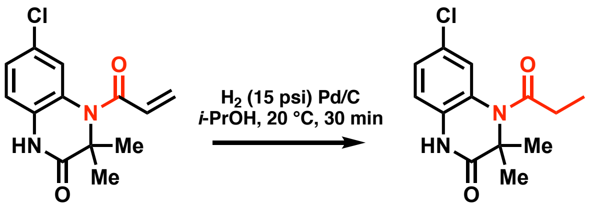


**EN1441** **CMZ139**

To a solution of Pd/C (25 mg, 10 wt. % loading) in 2-propanol (4 mL) under $N_2$, 6-chloro-3,3-dimethyl-4-prop-2-enoyl-1H-quinoxalin-2-one (50 mg, 188.89 μmol) was added. The reaction mixture was degassed and purged with $H_2$ three times and then stirred at 20 °C for 30 min under $H_2$ (15 psi) atmosphere. The reaction mixture was filtered, then concentrated *in vacuo*. Purification *via* prep-HPLC (column: Waters Xbridge BEH C18, 10 µm, 100 × 25 mm; mobile phase: [$H_2O$ (10 mM $NH_4HCO_3$) – acetonitrile]; gradient: 25% – 60% B over 8.0 min) gave 6-chloro-3,3-dimethyl-4-propanoyl-1H-quinoxalin-2-one (23.9 mg, 89.61 µmol, 47.4% yield, 100% purity) as a white solid. MS (ESI) *m/z* = 267.0 $[M+H]^+$; $\delta_H$ (400 MHz, DMSO-*d*$_6$) 10.75 (s, 1H), 7.29 - 7.16 (m, 2H), 6.98 (d, *J* = 8.6 Hz, 1H), 2.40 (q, *J* = 7.2 Hz, 2H), 1.45 (s, 6H), 1.03 - 0.95 (m, 3H). $\delta_C$ (101 MHz, DMSO-*d*$_6$) 176.02, 171.30, 129.77, 128.27, 125.78, 125.57, 123.42, 116.61, 60.30, 30.33, 22.98, 10.38. Calcd for $C_{13}H_{16}ClN_2O_2^+$ $[M+H]^+$ : 267.0895; found: 267.0902.

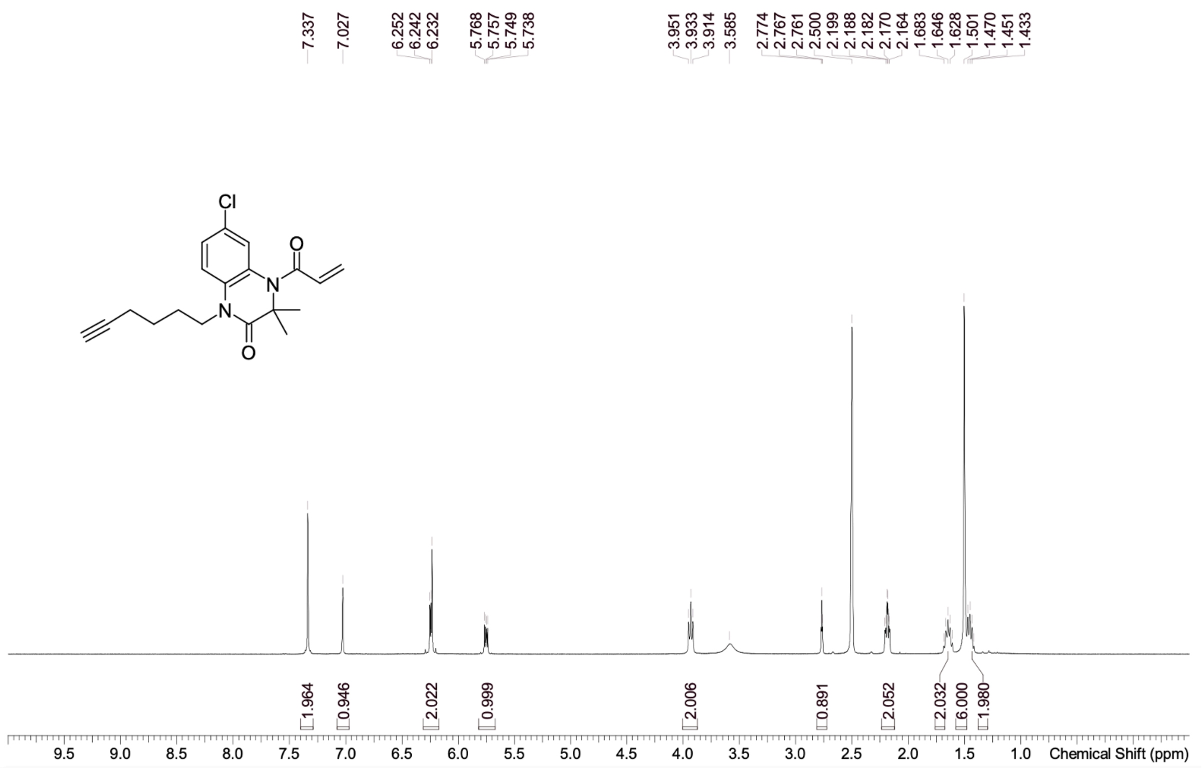
7.337
7.027
6.252
6.242
6.232
5.768
5.757
5.749
5.738
3.951
3.933
3.914
3.585
2.774
2.767
2.761
2.500
2.199
2.188
2.182
2.170
2.164
1.683
1.646
1.628
1.501
1.470
1.451
1.433
Cl
O
N
N
O
1.964
0.946
2.022
0.999
2.006
0.891
2.052
2.032
6.000
1.980
9.5 9.0 8.5 8.0 7.5 7.0 6.5 6.0 5.5 5.0 4.5 4.0 3.5 3.0 2.5 2.0 1.5 1.0
Chemical Shift (ppm)

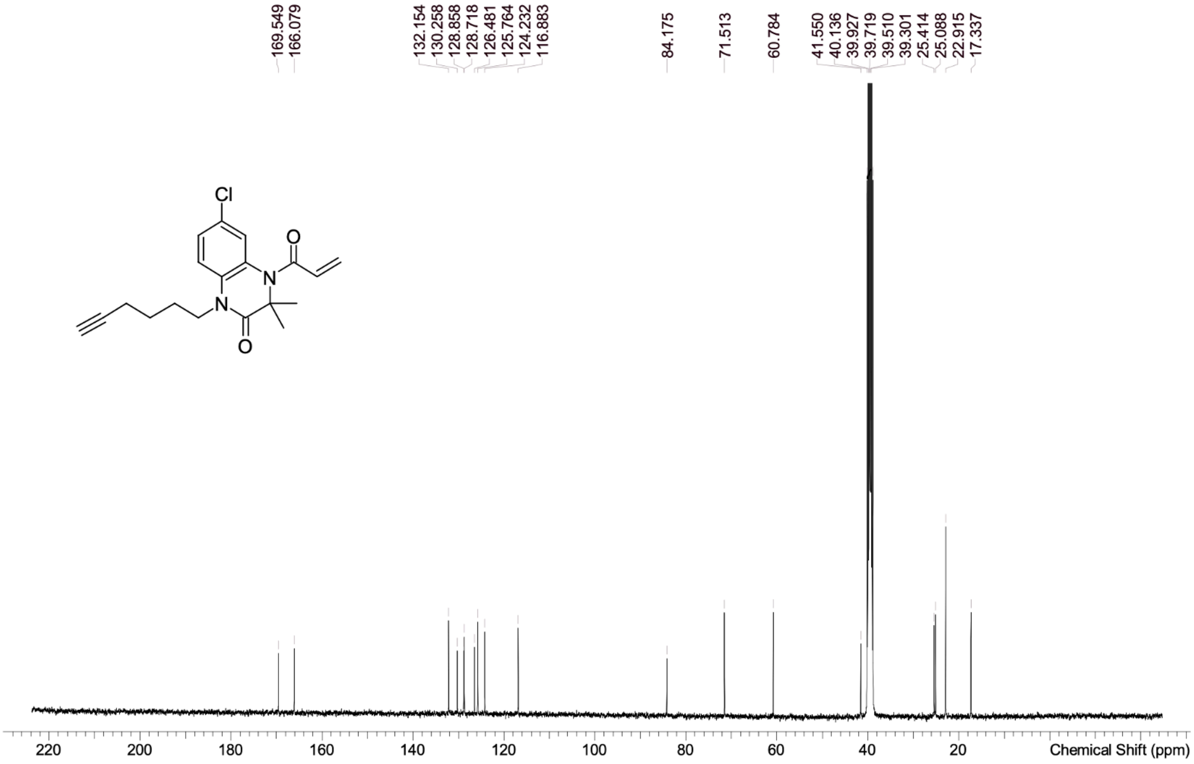
169.549
166.079
132.154
130.258
128.858
128.718
126.481
125.764
124.232
116.883
84.175
71.513
60.784
41.550
40.136
39.927
39.719
39.510
39.301
25.414
25.088
22.915
17.337
Cl
O
N
N
O
220 200 180 160 140 120 100 80 60 40 20
Chemical Shift (ppm)

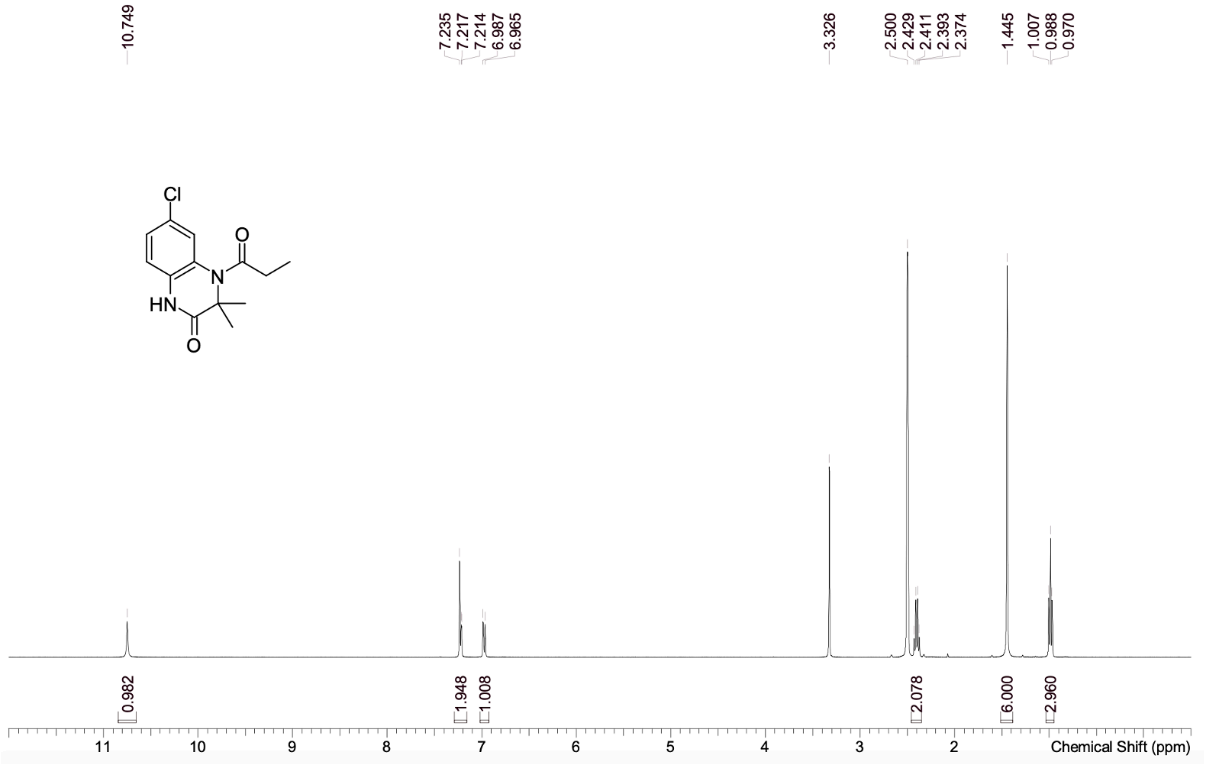
10.749
7.235
7.217
7.214
6.987
6.965
3.326
2.500
2.429
2.411
2.393
2.374
1.445
1.007
0.988
0.970
Cl
O
N
HN
O
0.982
1.948
1.008
2.078
6.000
2.960
11
10
9
8
7
6
5
4
3
2
Chemical Shift (ppm)

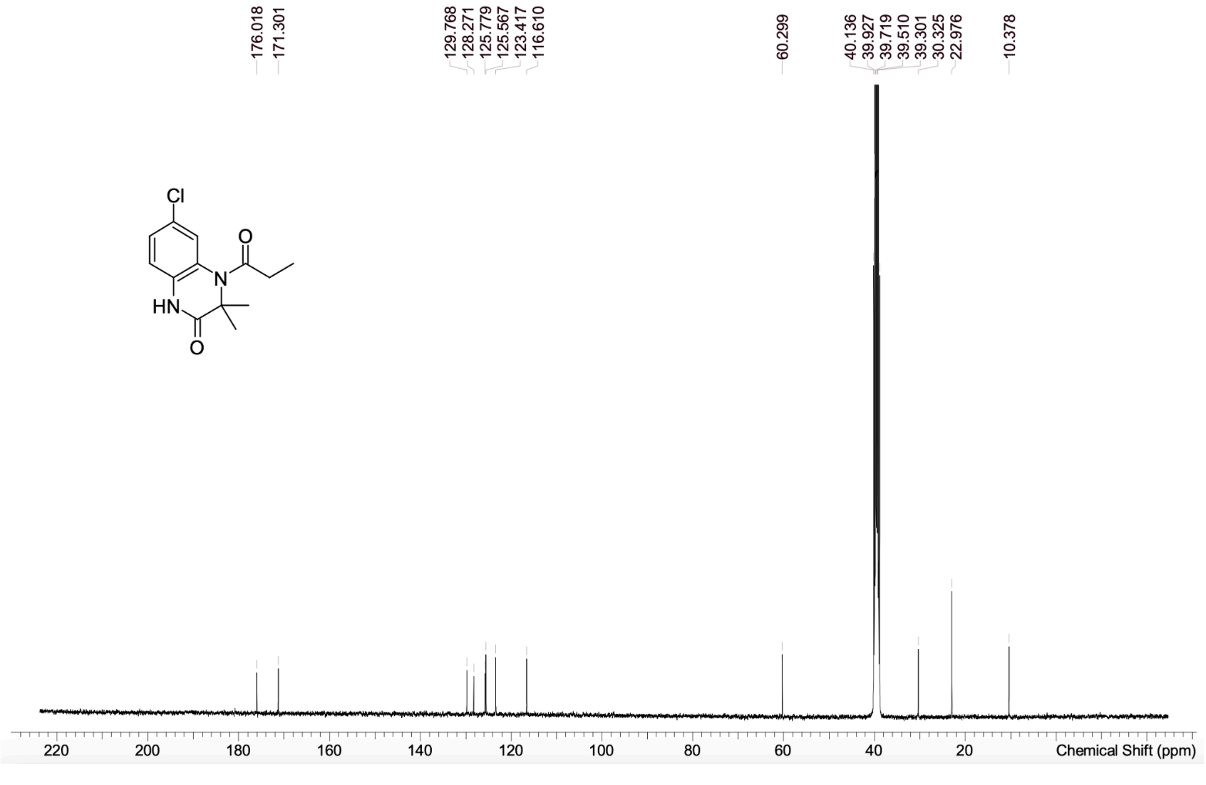
176.018
171.301
129.768
128.271
125.779
125.567
123.417
116.610
60.299
40.136
39.927
39.719
39.510
39.301
30.325
22.976
10.378
Cl
O
N
HN
O
220
200
180
160
140
120
100
80
60
40
20
Chemical Shift (ppm)
